# Combining Nanoscale Curvature and Polymer Osmotic Pressure for Efficient Giant Vesicle Assembly under Physiological Conditions

**DOI:** 10.1101/2025.02.14.638371

**Authors:** Alexis Cooper, Vignesha Vijayananda, Joseph Pazzi, Anand Bala Subramaniam

## Abstract

Thin film hydration methods are widely used to assemble giant unilamellar vesicles (GUVs), but their efficiency drops sharply in solutions of physiological ionic strength due to enhanced adhesion between stacked lipid bilayers, which suppresses bud and foam-like mesophase formation. Here, we introduce polymer-coated nanocellulose paper (PCP) as a nanostructured substrate that overcomes this limitation. Hydration of lipid films on PCP—termed PCP-assisted hydration— achieves high GUV yields across a broad temperature range (4–45 °C) using diverse soluble polymers, including ultralow-gelling agarose, hyaluronic acid, dextran, carrageenan, and polylysine. The nanoscale curvature of the cellulose fibers and the osmotic pressure generated by polymer dissolution act synergistically to promote membrane budding even under physiological salt conditions. The approach is scalable—supporting GUV production from millimeter-sized pieces to large-format sheets—and biocompatible, enabling encapsulation of complex biochemical systems such as cell-free expression mixtures and actin–fascin assemblies. PCP-assisted hydration thus provides a robust, versatile, and high-yielding platform for generating functional GUVs under physiological conditions.

## 1 Introduction

Self-assembly of lipid films in low salt aqueous solutions into surface-attached micrometer-sized buds or volume-spanning “foam”-like mesophases is a precursor to the formation of cell-sized giant unilamellar vesicles (GUVs) via thin film hydration methods.^[1,2]^ The GUV molar yield, *Y,* which is the amount of lipids in the membranes of the harvested GUVs relative to the amount of lipids deposited on the surface, reported as a percentage, allows comparison between different conditions of assembly.^[1,3]^ Qualitative categories classify the quantitative numbers into relative ranges, “negligible”, *Y* < 1 % “low”, 1 ≤ *Y* < 10%, “moderate”, 10 ≤ *Y* < 20 %, “high”, 20 ≤ *Y* < 45 % and “ultrahigh”, *Y* ≥ 45 %.

GUV yields obtained using thin film hydration methods, such as gentle hydration on glass slides, PAPYRUS (Paper-Abetted amPhiphile hYdRation in aqUeous Solutions), and electroformation, were moderate to high when performed using low salt solutions.^[4]^ However, in salty solutions, including those of physiological ionic strengths, the increased adhesion between the lamellar bilayers prevents the formation of buds on the surface or the formation of foam-like mesophases, resulting in negligible yields of GUVs.^[2]^ While GUVs encapsulating low salt solutions have myriad uses in biophysical and biological applications,^[5–10]^ the reconstitution of cellular proteins often requires a salty interior within the “physiological” range (50 – 150 mM of monovalent salt and 0 – 5 mM of divalent salt)^[11–18]^ marking a significant limitation to thin film hydration techniques. Depositing a dry film of partially soluble polymers, such as agarose of various gelling temperatures or polyvinyl alcohol (PVA), onto glass slides prior to lipid deposition allows the formation of GUVs in salty solutions.^[3,19,20]^ The polymer films partially dissolve and exert an osmotic pressure against the stacks of membranes, which overcomes the increased adhesion between the lipid bilayers.^[3,19]^ GUV yields mostly ranged from low to moderate. The highest measured yield, ∼ 17 %, was only achieved for low gelling temperature (LGT) agarose at 22 °C. ^[3]^ Agaroses of all other melting temperatures and polyvinyl alcohol (PVA) had yields below 10 %. LGT agarose at temperatures other than 22 °C had yields of < 5 %.^[3]^

Here, we report the fabrication of polymer-coated nanocellulose paper (PCP) which overcomes these limitations. We fabricate PCP using a wide variety of soluble macromolecules, including biocompatible and biomedically-relevant polymers such as hyaluronic acid, dextran, carrageenan, and polylysine. The PCP can be used to obtain GUV yields of > 15 % in salty solutions at a variety of temperatures and with membranes of varying compositions. We were inspired to fabricate PCP based on our earlier findings that the nanoscale curvature of nanocellulose fibers reduces the free energy of budding of lipid films^[4]^ in low salt solutions and that the osmotic pressure contributed by polymer dissolution on flat polymer-coated glass (PCG) can promote GUV bud formation in salty solutions^[3]^. We reasoned that coating nanocellulose fibers with dissolvable polymers would further enhance GUV yields in salty solutions by combining nanoscale curvature with osmotic pressure. We additionally find that patterns of dewetted polymers and lipid were smaller in scale on PCP compared to PCG. The apparent reduction in dewetting of the polymer on the surface of PCP further contributes to the enhanced yield of GUVs.

We demonstrate that the PCP-assisted hydration technique effectively prepares GUVs that encapsulate functional complex biochemical mixtures such as commercial cell-free protein expression systems and G-actin and the bundling protein fascin. The first achieves ∼100% deGFP expression from deGFP plasmids and the second forms single actin rings within the assembled GUVs. PCP-assisted hydration is highly adaptable, generating GUVs from PCP pieces as small as 3 mm in diameter with 12 µL of buffer to sheets as large as 279 mm × 178 mm in 200 mL of buffer. Under the latter conditions, the method produces approximately 6 × 10⁹ GUVs that encapsulate physiological saline.

## 2 Results and Discussion

### 2.1 Design of polymer-coated nanocellulose paper for assembly of GUVs in salty solutions

We were motivated to use nanocellulose paper as a substrate because its nanoscale cylindrical fibers facilitate the efficient assembly of GUVs in low salt solutions. Unlike flat surfaces, the nanoscale curvature of the fibers promotes spontaneous budding of conformal lipid bilayers, reflected by a negative free energy change. The free energy of forming a spherical bud of radius 𝑅_B_ from either cylindrical or disk-shaped bilayers is given by Equations (1) and (2) respectively^[3,4]^:

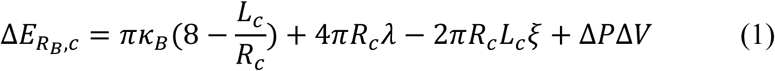

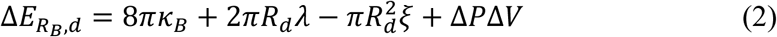

In these equations, 𝜅_B_ is the bending modulus, 𝜆 is the edge energy, 𝜉 is the adhesive potential (𝜉 < 0 for attraction), and Δ𝑃Δ𝑉 is the pressure–volume work. The geometric parameters are the cylinder radius, 𝑅_c_, cylinder length, 𝐿_c_, and disk radius, 𝑅_d_. The terms represent changes in bending, edge, adhesion, and pressure–volume energies, respectively.

Calculations show that budding from nanoscale cylindrical bilayers is strongly favored in low salt solutions but not in salty solutions. For example, for a bud of radius 𝑅_B_ = 141 nm, Δ𝐸_RB,c_ ≈ −4693 𝑘_B_𝑇, is negative in low salt solutions, but becomes positive, Δ𝐸_RB,c_ ≈ 1361 𝑘_B_𝑇, in high salt solutions. In contrast, bud formation from disk-shaped bilayers on flat surfaces is always less favorable, Δ𝐸_RB,d_ ≈ 4892 𝑘_B_𝑇 in low salt solutions and Δ𝐸_RB,d_ ≈ 10910 𝑘_B_𝑇 in salty solutions. These estimates use characteristic dimensions 𝑅_c_ = 20 nm, 𝐿_c_ = 2000 nm, 𝑅_d_ = 282 nm and characteristic parameters for DOPC bilayers, 𝜅_B_ = 8.5 × 10^-20^ J, 𝜆 = 1 × 10^-11^ J m^-1^, and 𝜉 = −1 × 10^-6^ J m^-2^ in low salt solutions and 𝜉 = −1 × 10^-4^ J m^-2^in salty solutions,^[21,22]^ with Δ𝑃 = 0.

Although budding becomes less favorable in salty solutions, the free energy cost is still lower for nanocellulose fibers than for flat substrates. Consistent with our earlier finding that polymer dissolution on glass coverslips can promote GUV formation in salty solutions by exerting an osmotic pressure, Δ𝑃, against the membrane, we reasoned that coating nanocellulose fibers with dissolvable polymers would further enhance GUV yields by combining nanoscale curvature with osmotic pressure.

### 2.2 Fabrication and characterization of polymer-coated nanocellulose paper

We design a series of experiments to compare the performance of PCP with polymer-coated glass slides (PCG). We expect that due to the nanoscale curvature of the fibers, controlling the surface concentration of polymer will be important to preserve the effects of curvature while having sufficient polymer to promote budding. Previous studies have reported a range of agarose and PVA concentrations on glass substrates (Supporting Text and Table S1). Thus, we systematically varied both the surface concentration of the polymer and the substrate identity to assess their combined influence on GUV assembly. Scanning electron microscopy (SEM) confirmed that the nanocellulose paper consists of a network of randomly entangled, polydisperse cylindrical fibers (Figure 1a). The glass slide in contrast appears smooth and featureless (Figure 1b). We use the polymer ultralow gelling temperature (ULGT) agarose to create PCP and PCG substrates. We prepared ULGT agarose PCP and PCG with polymer nominal surface concentration (polymer-NSC) of 0, 0.05, 0.5, 1.5, and 5.2 nmol/cm^2^. We then deposit 10 µg of a lipid mixture containing 96.5 mol % 1,2-dioleoyl-*sn*-glycero-3-phosphocholine (DOPC), 3 mol % of 1,2-dioleoyl-*sn*-glycero-3-phosphoethanolamine-*N*-[methoxy(polyethylene glycol)-2000] (ammonium salt) (PEG2000-DSPE) and 0.5 mol % TopFluor^™^ Cholesterol (TopFluor-Chol). We allow the assembly to proceed for 1 hour in phosphate-buffered saline (PBS) + 100 mM sucrose. PBS has a composition of 137 mM sodium chloride, 2.7 mM potassium chloride, 8 mM sodium phosphate dibasic, 2 mM potassium phosphate monobasic, and is at a pH of 7.4. PBS is osmotically balanced with cells and is a widely used buffer in biological research. PEG2000-DSPE provides steric repulsion that inhibits the aggregation of GUVs in the salty buffer. TopFluor-Chol is a fluorescent sterol that allows visualization of the lipid membranes through confocal fluorescence microscopy. The sucrose allowed us to obtain a density gradient between the GUVs and the outer solution for sedimentation and imaging.

**Figure 1.**
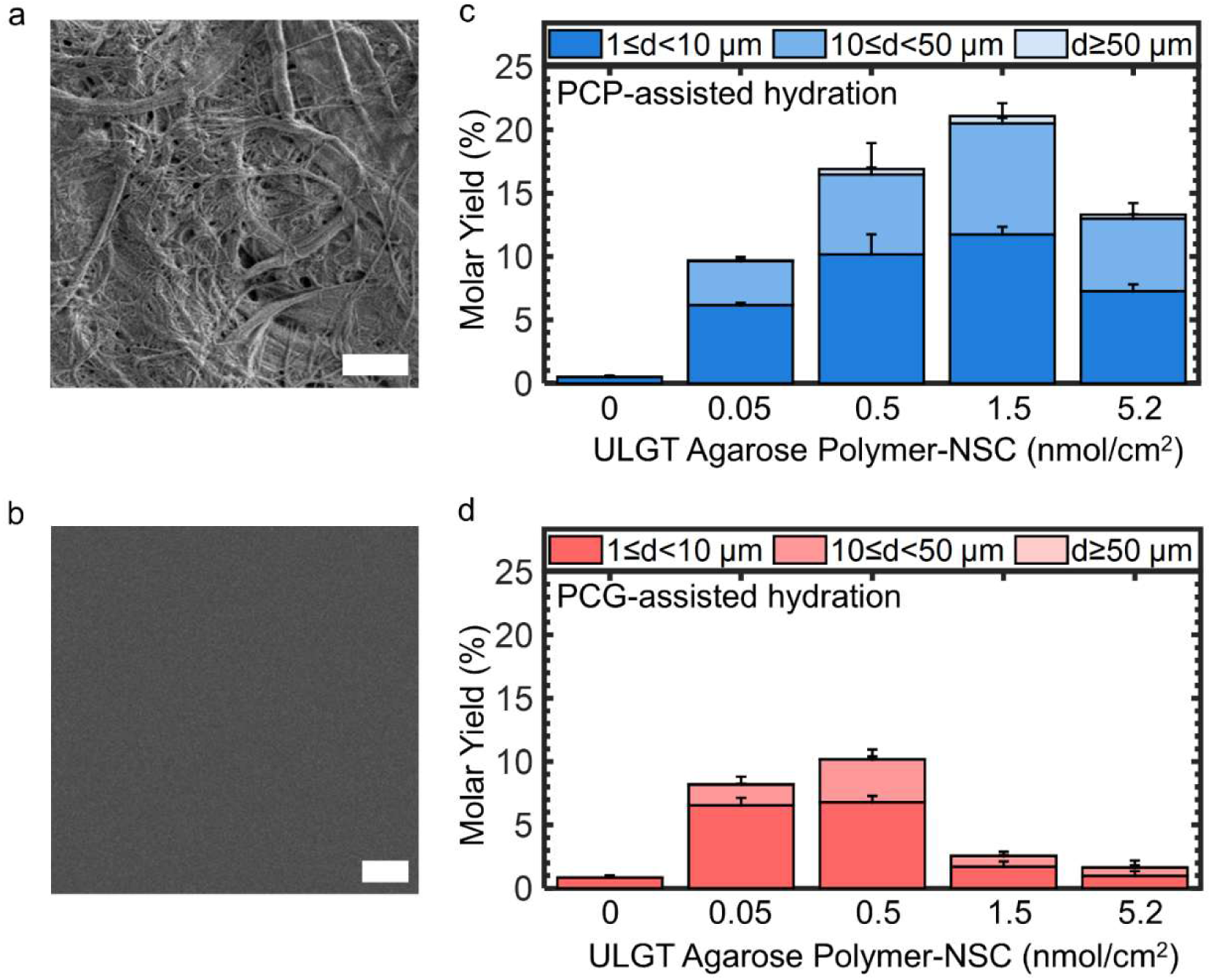
Surface morphology of the substrates and yields of GUVs obtained using PCP- and PCG-assisted hydration at various ULGT agarose polymer-NSCs. a) SEM image of nanocellulose paper. b) SEM image of a glass slide. c) Stacked bar plots of GUV yields from PCP-assisted hydration. d) Stacked bar plots of GUV yields from PCG-assisted hydration. The stacks show the percentage of the molar yield that is comprised of GUVs with the diameters listed in the legend. The bar is an average of N=3 independent repeats. The error bars are one standard deviation from the mean. Scale bars are 2 µm.

After 1 hour of incubation, we use a micropipette to aspirate the solution. The fluid shear due to aspiration detaches the buds from the surface, which then self-close to form GUVs (See Experimental Section for details). Similar to previous results with PAPYRUS and gentle hydration on glass slides, we confirmed that the majority, if not all, of the buds are harvested from the polymer-coated surfaces (Figure S1). We use a custom MATLAB routine to obtain the distribution of diameters, counts, and the molar yield of the GUVs. The routine excludes multilamellar vesicles, bright lipid aggregates, and vesicles < 1 µm, which we term non-GUV structures.^[3,4]^ Figure 1c and 1d are stacked bar plots showing the evolution of the molar yields as a function of polymer-NSC for ULGT agarose PCP and PCG respectively. We show representative images of the harvested GUVs and histograms of the diameters in Figure S2 and S3.

In the absence of ULGT agarose, the molar yield was negligible, < 1 %, for both substrates. PCP produced a higher yield of GUVs than PCG for all the polymer-NSCs that we tested. Both these results are consistent with the free energy change expected for budding in salty solutions. Both surfaces show a peaked distribution (Figure 1c, d). The location of the peak and the yield at the peak differed between PCP and PCG substrates. The maximum yield obtained using ULGT agarose PCG-assisted hydration was 10.2 ± 0.8 % at a polymer-NSC of 0.5 nmol/cm^2^. The maximum yield obtained using ULGT agarose PCP was double that of the ULGT agarose PCG, 21.1 ± 1.0 %, at a polymer-NSC of 1.5 nmol/cm^2^. We term the polymer-NSC that produces the maximum yield as the “optimal” polymer-NSC.

The differences in yield have practical implications. The number of GUVs with diameters > 10 µm scales linearly with molar yield (Figure 2a,b, Pearson’s R = 0.9933 and 0.9607 respectively). Thus, substrates that produce higher molar yields not only improve overall GUV formation but also favors the generation of larger GUVs compared to substrates that produce lower molar yields. Large GUVs are advantageous for biophysical experiments and imaging. Additionally, a fortuitous observation was that the samples obtained using PCP-assisted hydration had consistently lower amounts of non-GUV structures in the solution compared to the samples obtained using PCG-assisted hydration (Figure 2c,d, white arrows, see Figure S2 for additional images). For example, even when the molar yields of GUVs were similar, ∼ 10 %, the percentage of non-GUV structures from PCG-assisted hydration was more than double that of PCP-assisted hydration, 19.4 ± 0.6 % versus 8.5 ± 0.3 % respectively. See Figure S3 for the percentage of non-GUV structures for the other polymer-NSCs.

**Figure 2.**
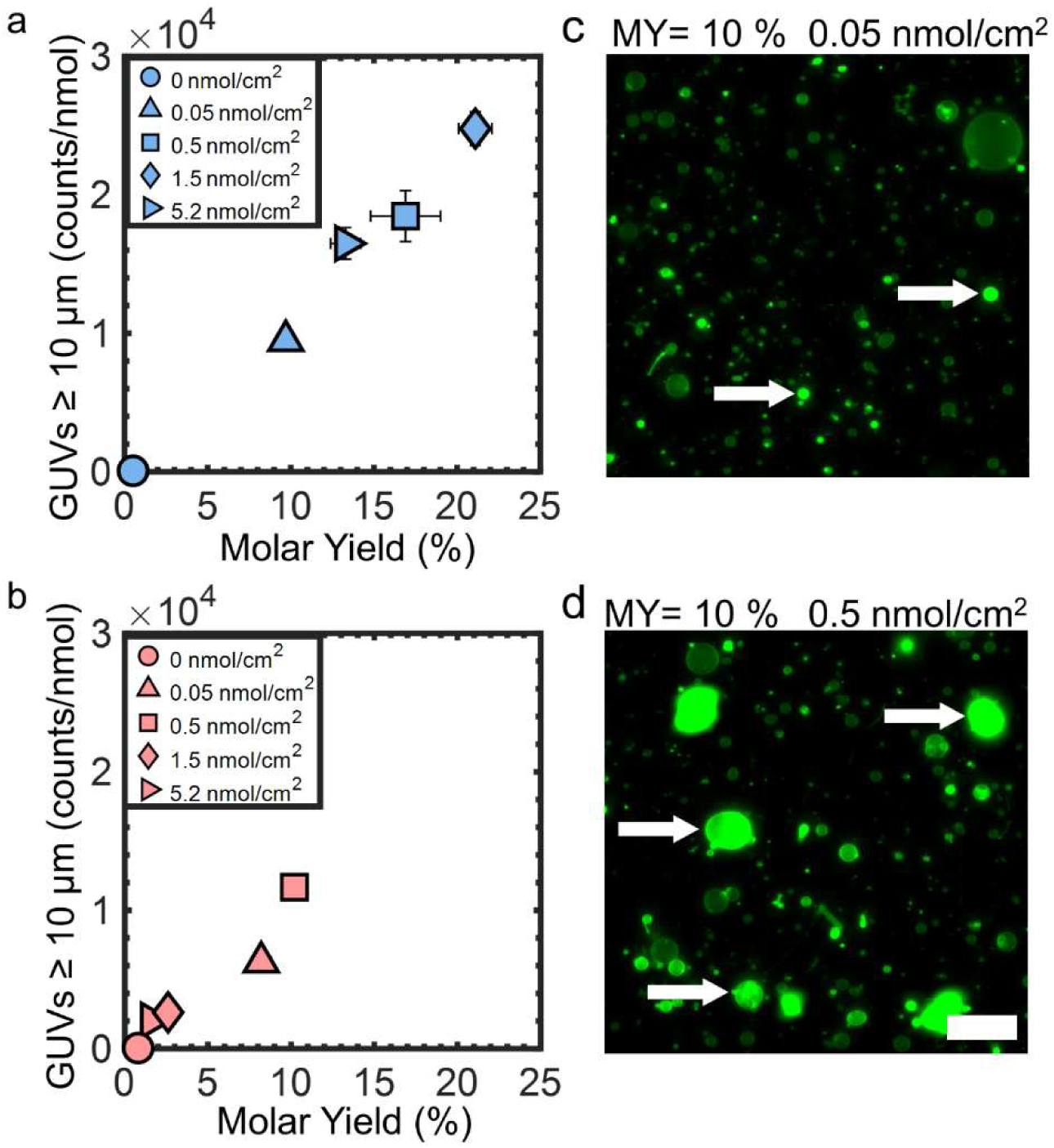
Relationship between the number of GUVs with diameters ≥ 10 µm and the molar yield and number of non-GUV structures. a) Scatter plot of the counts of GUVs with diameters ≥ 10 µm versus the total molar yield obtained using PCP-assisted hydration. b) Scatter plot of the counts of GUVs with diameters ≥ 10 µm versus the total molar yield obtained using PCG-assisted hydration. c-d) Representative single-plane confocal microscopy images of harvested objects obtained from PCP-and PCG-assisted hydration. c) ULGT agarose PCP with a polymer-NSC of 0.05 nmol/cm^2^. d) ULGT agarose PCG with a polymer-NSC of 0.5 nmol/cm^2^. GUVs appear as circles with uniform membrane fluorescence intensity. The white arrows point to examples of non-GUV structures which have high fluorescence intensity and irregular shapes. Scale bar is 50 µm.

We perform experiments using two other commercially available substrates, frosted glass slides and nitrocellulose membranes, to determine if surface roughness or porosity alone can recapitulate the results of PCP. Figure 3a and 3b show SEM images of the frosted glass slides and nitrocellulose membranes respectively. Large lipid-polymer pseudobuds, characterized by the presence of large dark non-fluorescent areas at the base of the buds, formed on polymer-coated frosted glass (Figure 3c, white arrow). These pseudobuds are non-productive for forming GUVs. ^[3]^ Sparse buds formed on the surface of the polymer-coated nitrocellulose membranes (Figure 3d).

**Figure 3.**
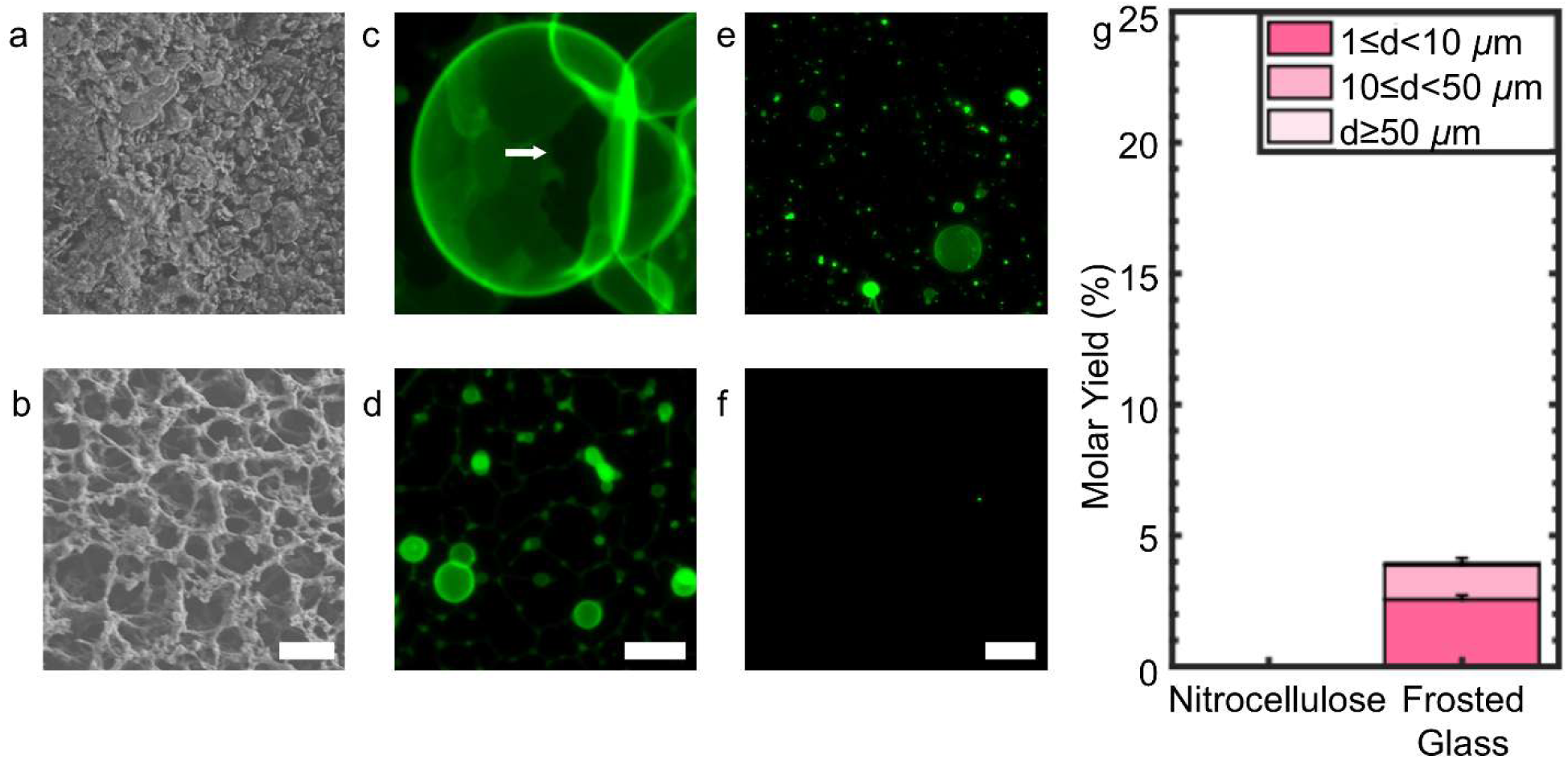
Surface morphology of the substrates and yields of GUVs obtained from polymer-coated frosted glass slides and nitrocellulose membranes. a) SEM image of a frosted glass slide. b) SEM image of a nitrocellulose membrane. c,d) Two-dimensional projection of *z*-slices from *z* = 0 to 100 µm from the surface using the “sum slices” algorithm of the surface morphology of DOPC films on the substrates that are coated with ULGT agarose at a polymer-NSC of 1.5 nmol/cm^2^. c) The frosted glass slide and d) the nitrocellulose membrane. The white arrow highlights a dewetted region of lipid-polymer. e,f) Representative single plane confocal images of the objects that are harvested from the polymer-coated surfaces. e) The frosted glass slide and f) the nitrocellulose membrane. g) Stacked bar plots of molar yields obtained from each surface. The yield was not measurable for the ULGT agarose-coated nitrocellulose membrane. The stacks show the percentage of the molar yield that is comprised of GUVs with the diameters listed in the legend. The bar is an average of N=3 independent repeats. The error bars are one standard deviation from the mean. Scale bar in a-b) is 2 µm, c-d) 10 µm, and e-f) 50 µm.

Few GUVs were recovered from the polymer-coated frosted glass slides, and there were no GUVs from the polymer-coated nitrocellulose membrane (Figure 3e-f). The low yield of GUVs on polymer-coated frosted glass, and the lack of GUVs from polymer-coated nitrocellulose membranes, indicate that the entangled cellulose nanofibers of nanocellulose paper, and not merely surface roughness, is important for the superior performance of PCP (Figure 3g, see the histograms of diameters in Figure S4).

### 2.3 Dewetting is limited to smaller scales on polymer-coated nanocellulose paper

We next examined the lipid films on the PCP and PCG substrates one hour after hydration, that is before we harvest the buds, by capturing high-resolution confocal Z-stacks. The left panels, upper right panels, and lower right panels of Figure 4 show two-dimensional orthogonal *x-z* reconstructions, *x-y* projections of the entire bud layer, and *x-y* projections of the first 18.62 μm near the surfaces respectively. The pixel intensity histograms of the orthogonal reconstructions were equalized to show bright and dark regions in the image. See Figure S5 for the images without histogram equalization. Buds appear as stacked, size-stratified layers extending up to 60 µm from the surface in the orthogonal reconstruction at the optimal polymer-NSC on PCP (Figure 4a). Projections of the entire bud layer showed a dense layer of GUV buds. Additionally, the lipid layer on the surface of PCP showed few dark non-fluorescent regions, and the non-fluorescent regions that were present were small (Figure 4a, white arrows), indicative of few pseudobuds. In contrast, PCG at the optimal polymer-NSC showed a single monolayer of buds with most of the buds showing large non-fluorescent regions on the surface (Figure 4b, white arrows), indicative of many pseudobuds. Thus, we suggest that in addition to promoting budding, the polymer-coated nanocellulose paper appears to restrict dewetting to smaller length scales compared to the polymer-coated glass slides.

**Figure 4.**
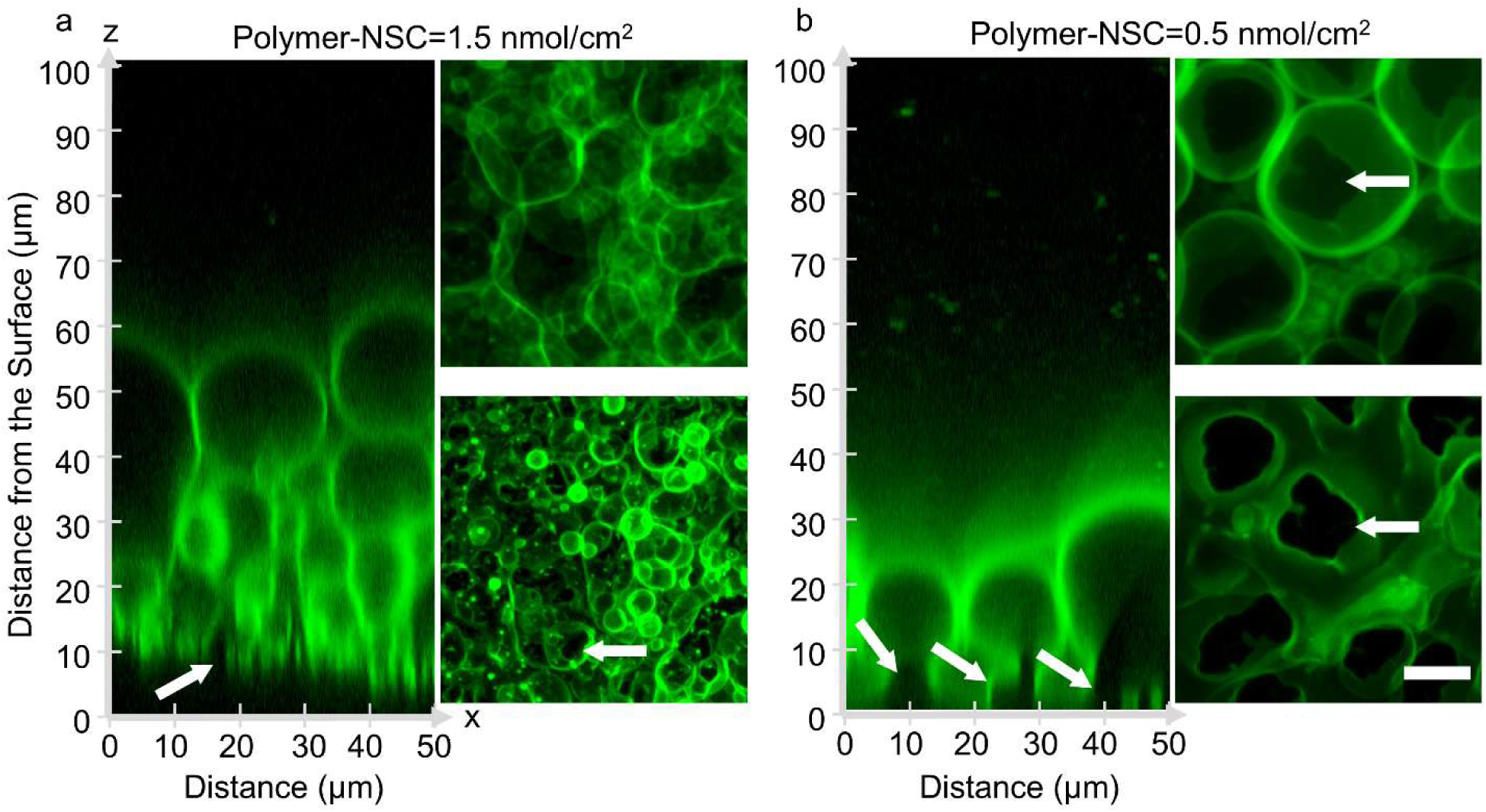
Configuration of the hydrated lipid films on ULGT agarose PCP and PCG at their optimal polymer-NSCs. a) PCP. b) PCG. The left panels show orthogonal *x*–*z* planes of confocal Z-stacks of the lipid films after 1 hr of hydration. The top right panels are two-dimensional projections of *z*-slices from 0 to 99.87 µm from the surface using the *‘sum slices’* algorithm. The bottom right panels are two-dimensional projections of *z*-slices from 0 to 18.62 µm using the “*maximum intensity projection”* algorithm. The white arrows point to examples of regions of dewetted lipid-polymer on the surfaces. The scale bar is 10 µm.

### 2.4 PCP-assisted hydration is compatible with diverse conditions of assembly

Having shown that PCP demonstrates superior performance compared to PCG at room temperature, we next test the performance of PCP-assisted hydration at different temperatures. Previously, we found that assembly at temperatures of ≥ 37 °C resulted in a greatly reduced yield of GUVs using LGT and ULGT agarose-coated glass coverslips.^[3]^ Higher than room temperature hydration and incubation of the lipid film is often required since bud formation occurs efficiently when the membrane is in a fluid phase, that is when the membrane is above the chain-melting transition of the constituent lipids. For membranes composed of mixtures of lipids, the assembly temperature must be above the lipid with the highest chain melting temperature.^[23]^ In contrast, hydration and incubation below room temperature, typically at 4 °C, with membranes composed of lipids with low chain melting temperatures, prevents premature reaction when constructing synthetic cells.^[11,14,15]^ For future cold-chain biomedical applications, assembly and storage of GUVs at low temperature could be useful for reducing the rate of activity of degradative enzymes, the rate of growth of microorganisms, and the rate of intrinsic degradation of biomolecules.^[24,25]^

We perform PCP-assisted hydration using a canonical ternary lipid mixture that exhibits “raft” phases at room temperature. The mixture consists of DOPC:1,2-dipalmitoyl-*sn*-glycero-3-phosphocholine (DPPC):Cholesterol:DSPE-PEG2000:1,2-dioleoyl-*sn*-glycero-3-phosphoethanolamine-*N*-(lissamine rhodamine B sulfonyl) (ammonium salt) (Rhod-DOPE): TopFluor-Chol at a mol % of 36:33:27.5:3:0.25:0.25. The membranes of the GUVs are expected to phase separate into equal area fractions of a DOPC-enriched liquid disordered (L_d_) phase and a DPPC- and cholesterol-enriched liquid ordered (L_o_) phase when quenched to room temperature.^[23]^

The Rhod-DOPE partitions into the L_d_ phase while the TopFluor-Chol partitions into the L_o_ phase, allowing visualization of these phases in the membranes of the GUVs. The chain melting temperature of DPPC is 41.5 °C ^[26]^ and thus we performed PCP-assisted hydration at 45 °C.^[23]^ The yield of the phase-separating GUVs was 17.0 ± 2.7 % after 1 hour which was statistically indistinguishable from the yield of single phase DOPC GUVs obtained using PCP-assisted hydration at 22 °C (*p* = 0.118) (Figure 5a, see Figure S6 for the histogram of diameters). Furthermore, images of the phase-separating GUVs showed equal area fractions of L_d_ and L_o_ phases (Figure 5b). To quantify the phase separation, we measured the diameters, *d*, and the height of the spherical caps, *h* that compose the L_d_ phase using FIJI. We find that the fraction of the L_d_ phase, 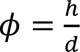, was 0.48 ± 0.06, of n = 75 (Figure 5c). This result shows that PCP-assisted hydration produces GUVs with compositions that largely reflect the composition of the lipid stock.^[27]^

**Figure 5.**
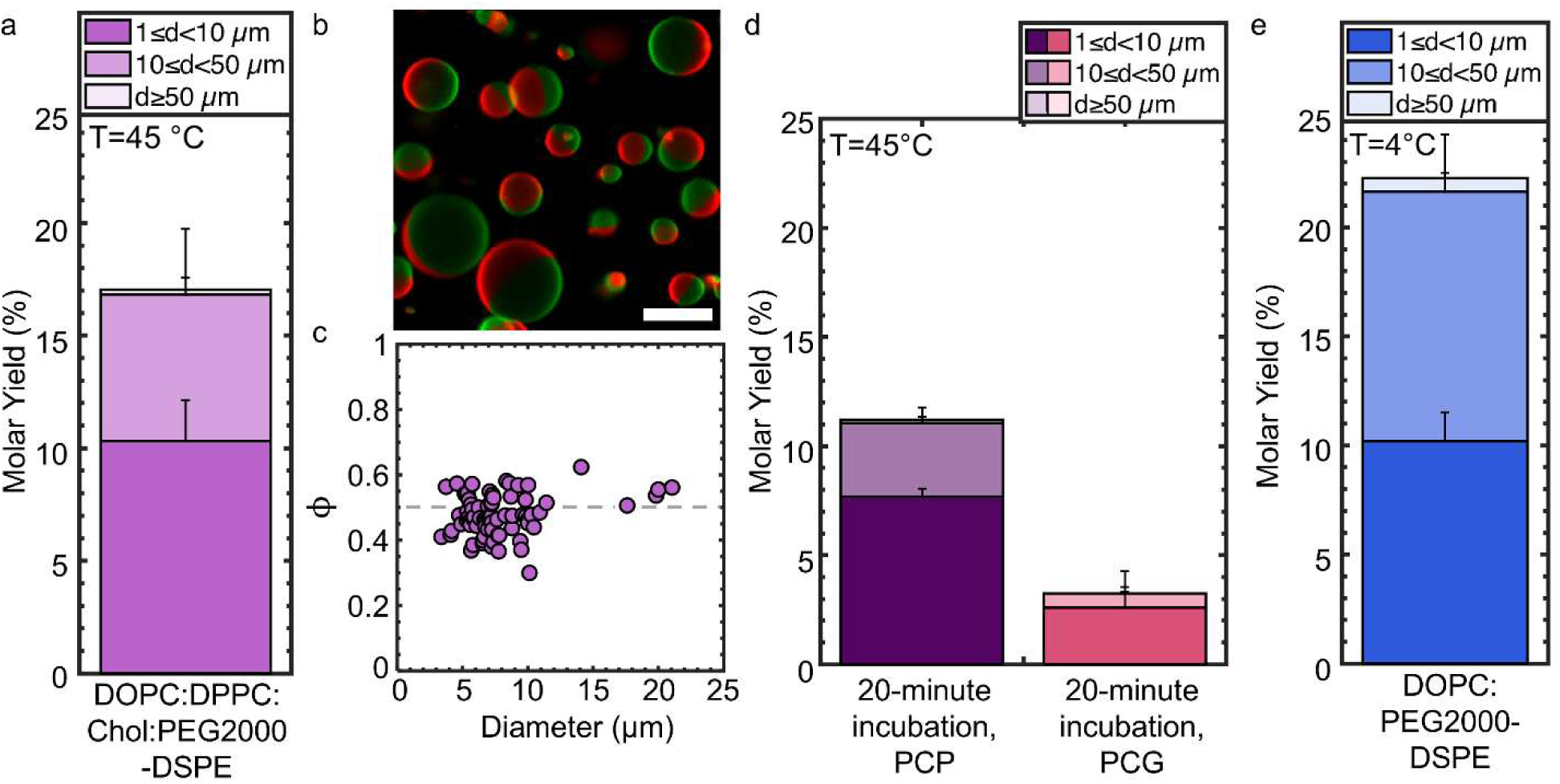
Molar yields of GUVs assembled at different temperatures using PCP- and PCG-assisted hydration. All substrates were prepared with ULGT agarose at the optimal polymer-NSC. a) Molar yield of DOPC:DPPC:Chol:PEG2000-DSPE phase-separating GUVs obtained using PCP-assisted hydration after 1 hour of incubation at 45 °C b) Representative dual channel confocal image of the GUVs at 22 °C showing phase separation. The L_d_ phase is false-colored red. The L_o_ phase is false colored green. c) Scatter plot of the fraction of the L_d_ phase, 𝜙, relative to the GUV diameter. n = 75 GUVs. d) Molar yield of phase-separating GUVs obtained using PCP-assisted hydration and PCG-assisted hydration after 20 minutes of incubation at 45 °C. e) Molar yield of DOPC:PEG2000-DSPE GUVs assembled at 4 °C using PCP-assisted hydration. The stacks in the bar plots show the percentage of the molar yield that is comprised of GUVs with the diameters listed in the legend. Each bar is the average of N=3 independent repeats. The error bars are one standard deviation from the mean. Scale bar in c) is 10 µm.

Although higher than room temperature assembly is required for membranes composed of high transition temperatures, high temperatures lead to time-dependent denaturation of proteins.^[28]^ It was previously shown that a shorter 20-minute incubation time preserves the functionality of membrane proteins which otherwise denatured when incubated for longer periods at 45 °C.^[28]^ We thus investigated the effect of incubation time on PCP-assisted hydration. The yield of the phase-separating GUVs obtained using PCP-assisted hydration was 11.2 ± 0.6 % after 20 minutes of incubation. This yield was more than 3× higher than the 3.2 ± 1.0 % yield of phase-separating GUVs that was obtained using PCG-assisted hydration under similar conditions (Figure 5d, see Figure S6 for the histogram of diameters).

To investigate assembly at low temperatures, we perform PCP-assisted hydration at 4 °C using DOPC. We use DOPC since its chain melting transition temperature is – 17 °C.^[29]^ We obtained a yield of 22.3 ± 2.0 % (Figure 5e). This yield was statistically indistinguishable from the yield obtained using PCP-assisted hydration at room temperature (*p* = 0.499). We show a representative image and histograms of the diameter distributions in Figure S7. In contrast, use of the LGT agarose PCP resulted in a significant reduction in yield from 14.8 ± 0.5 % at 22 °C to 8.6 ± 1.0 % at 4 °C (Table S2). We discuss in Supporting Text that the differences in solubility between ULGT agarose and LGT agarose at different temperatures could explain these observations.

We conclude that PCP-assisted hydration supports the efficient assembly of GUVs at a variety of temperatures and incubation times.

### 2.5 Polymer-coated nanocellulose paper can be fabricated using a wide range of macromolecules and polymers

Assembly of GUVs using assisting compounds necessarily leads to the incorporation of the compounds in the resulting GUVs.^[3,30,31]^ Since the incorporation of polymers is a necessary feature of the technique, we explored if there are limits to the identity of the polymer. Extant reports use partially-soluble^[19,20]^ or cross-linked polymers^[32,33]^ on smooth glass surfaces, overwhelmingly ultra-low gelling temperature agarose or PVA. To provide the greatest contrast with existing reports and to enable future applications of GUVs in the body, we chose from a selection of highly soluble polymers with known biomedical applications^[34–43]^. We use three polysaccharides, hyaluronic acid, dextran, and carrageenan, and a polypeptide, poly-D-lysine. Hyaluronic acid is an anionic polymer used in wound healing, in tissue engineering, and in cosmetics.^[35]^ Dextran is a clinically-approved plasma expander^[36]^ and a contrast agent for magnetic resonance imaging.^[37]^ When chemically cross-linked, dextran hydrogels are used in tissue engineering applications.^[38,39]^ Carrageenan is a gel-forming polymer that is used as a scaffold in tissue engineering,^[42]^ in drug delivery applications,^[41,43]^ and in food to modify viscosity and texture.^[41]^ Polylysine is used extensively as a promoter for cell adhesion in cell culture.^[40]^ We used hyaluronic acid of two different molecular weights (MW), 8-15 kDa and 70-120 kDa, dextran with a MW of 100 kDa, carrageenan with a MW of 521 kDa, and poly-D-lysine with a MW of 70-150 kDa. We also tested fragmented salmon DNA that was ∼ 587-837 base pairs with a MW of 274 kDa and bovine serum albumin (BSA) with a MW of 66 kDa. These two compounds are not used for biomedical applications but have different macromolecular and chemical characteristics compared to the other polymers that we tested. Salmon DNA is a nucleic acid and BSA is a globular protein. We tested various polymer-NSCs, lipid compositions, and assembly temperatures. We report in Table 1 the optimal polymer-NSC, molar yield, the GUV counts normalized to the mass of lipid that was deposited on the surface, and the percentage of non-GUV structures. We present additional results and discussion from our screen of various polymer types, polymer-NSC, and temperatures in Supporting Text and Tables S3-S5.

**Table 1.** Polymer type, optimal polymer-NSC, lipid composition, molar yields, GUV counts per µg of lipid, and the percent non-GUV objects. Each value is reported as a mean ± standard deviation of N=3 independent repeats.

| Polymer | Optimal Polymer-NSC (nmol/cm<sup>2</sup>) | Lipid Composition | Molar Yield (%) | GUV Counts ($\times 10^5$ GUVs/$\mu\text{g}$) | Percent non-GUVs (%) |
| --- | --- | --- | --- | --- | --- |
| **4°C** |  |  |  |  |  |
| Agarose, ULGT*<br>120 kDa | 1.5 | DOPC:PEG2000-DSPE | 22.3 ± 2.0 | 5.4 ± 0.6 | 7.2 ± 0.3 |
| Dextran,<br>100 kDa | 1.9 | DOPC:PEG2000-DSPE | 17.7 ± 0.7 | 8.5 ± 0.8 | 10.1 ± 0.4 |
| Hyaluronic Acid,<br>8-15 kDa | 23 | DOPC:PEG2000-DSPE | 20.4 ± 0.9 | 10.8 ± 0.6 | 8.1 ± 0.2 |
| Hyaluronic Acid,<br>70-120 kDa | 2.7 | DOPC:PEG2000-DSPE | 12.6 ± 2.7 | 5.2 ± 1.6 | 9.8 ± 1.3 |
| **22 °C** |  |  |  |  |  |
| Agarose, ULGT*<br>120 kDa | 1.5 | DOPC:PEG2000-DSPE | 21.1 ± 1.0 | 5.8 ± 0.5 | 8.3 ± 0.5 |
| Agarose, ULGT*<br>120 kDa | 1.5 | DOPC:DGS-NTA(Ni):PEG2000-DSPE | 18.1 ± 0.7 | 5.7 ± 0.5 | 5.3 ± 0.5 |
| Bovine Serum<br>Albumin | 2.8 | DOPC:PEG2000-DSPE | 6.1 ± 0.5 | 3.4 ± 0.4 | 10.6 ± 0.7 |
| Carrageenan,<br>521 kDa | 1.2 | DOPC:PEG2000-DSPE | 16.4 ± 2.0 | 6.3 ± 1.2 | 7.0 ± 2.0 |
| Dextran,<br>100 kDa | 4.3 | DOPC:PEG2000-DSPE | 15.0 ± 0.3 | 5.8 ± 0.1 | 11.3 ± 0.1 |
| DNA, Single<br>stranded from<br>salmon testes | 2.3 | DOPC:PEG2000-DSPE | 6.7 ± 1.2 | 6.1 ± 1.4 | 9.7 ± 0.3 |
| Hyaluronic Acid,<br>70-120 kDa | 2.7 | DOPC:PEG2000-DSPE | 15.5 ± 2.0 | 5.2 ± 1.6 | 9.8 ± 1.3 |
| Hyaluronic Acid,<br>8-15 kDa | 23 | DOPC:PEG2000-DSPE | 22.0 ± 2.5 | 10.1 ± 0.7 | 10.8 ± 0.1 |
| Poly-D-Lysine,<br>70-150 kDa | 6.2 | DOPC:PEG2000-DSPE | 18.6 ± 3.0 | 6.8 ± 1.4 | 9.3 ± 1.9 |
| **45°C** |  |  |  |  |  |
| Agarose, ULGT*<br>120 kDa | 1.5 | DOPC:DPPC:<br>Chol:PEG<br>2000-DSPE | 17.0 ± 2.7 | 5.0 ± 1.0 | 7.0 ± 0.03 |
| Hyaluronic Acid, 8-15 kDa | 23 | DOPC:DPPC:<br>Chol:PEG<br>2000-DSPE | 17.5 ± 2.2 | 8.4 ± 1.6 | 15.8 ± 1.1 |

\*Data from ULGT agarose PCP-assisted hydration reproduced from Fig. 1 and 5 for ease of comparison

We find that most of these molecules when used to prepare PCP substrates support the formation of GUVs at moderate to high yields via PCP-assisted hydration, 13 - 22 %, at the various temperatures of assembly that we tested. Use of hyaluronic acid PCP to assemble GUVs results in high yields of 20.4 ± 0.9 %, 22.0 ± 2.5 %, and 17.5 ± 2.2 % at 4 °C, 22 °C, and 45 °C, respectively. Use of dextran PCP to assemble GUVs resulted in yields of 17.7 ± 0.7 % and 15.0 ± 0.3 % at 4 °C and 22 °C. Use of poly-D-lysine PCP to assemble GUVs resulted in a high yield of 18.6 ± 3.0 % at 22 °C. Use of nanocellulose paper as a substrate was essential for obtaining high yields of GUVs using these highly soluble polymers. The use of glass as a substrate resulted in massive dewetting of the lipid-coated polymer films and a negligible yield of GUVs (Figure S8). Although DNA PCP and BSA PCP resulted in higher yields of GUVs compared to using bare nanocellulose paper, the yields were lower than the other molecules we tested, at 6.7 ± 1.2 % and 6.1 ± 0.5 %, respectively.

We find no general correlation between the molecular weight and optimal polymer-NSC with the yields of GUVs obtained using PCP-assisted hydration (Table 1). This observation is consistent with more than one process, polymer interactions with the membrane, dewetting, solubility, and the rate of dissolution, affecting the yield.^[3]^ Thus, the polymer-NSC must be optimized for each novel macromolecule when fabricating PCP for use in PCP-assisted hydration.

### 2.6 PCP-assisted hydration produces higher yields and larger GUVs compared to electroformation in salty solutions

To the best of our knowledge, other than gel-assisted hydration (PCG-assisted hydration in the nomenclature used in this paper), ^[18–20]^ only high-frequency (HF) electroformation is reported to be able to produce high yields GUVs in salty solutions among thin film hydration techniques. ^[44,45]^ Studies of HF electroformation use membranes that do not contain lipids with PEGylated headgroups. ^[44,45]^ We compare the lipid composition that is reported in the literature with the mixture that we use in this work by using lipid mixtures consisting of DOPC that excluded the PEG2000-DSPE lipid (0 mol % PEG2000-DSPE). We use the high-frequency (HF) protocol ^[44,45]^ and a low-frequency (LF) protocol, which is widely used to assemble GUVs in low salt solutions.^[1,4,46,47]^ For the HF protocol, we applied a sinusoidal AC field at a frequency of 500 Hz and at variable field strengths. We use a field strength of 0.106 V/mm peak-to-peak for 5 minutes, 0.940 V/mm peak-to-peak for 20 minutes, and 2.61 V/mm for 90 minutes, and then imaged the configuration of the lipid film on the surface or harvested the buds. For the LF protocol, we applied a sinusoidal AC field at a frequency of 20 Hz for 2 hours at a constant field strength of 1.5 V/mm peak-to-peak. The 0 mol % PEG2000-DSPE composition had few buds, and most of them were aggregated on the surface (Figure 6a). The suspension showed a few aggregated GUVs (Figure 6b). This result is expected as PBS contains 137 mM of sodium chloride. Charges are effectively screened, as the Debye screening length is 0.75 nm at this electrolyte concentration, and zwitterionic membranes adhere. Since most of the GUVs were aggregated as clusters, it was not possible to obtain a quantitative molar yield. In contrast, for the mixture containing 3 mol % PEG2000-DSPE, there were isolated spherical buds (Figure 6c) and the harvested sample contained colloidally-stable isolated GUVs (Figure 6d). The molar yield was 5.9 ± 1.2 % for the LF protocol and 10.0 ± 1.9 % for the HF protocol (Figure 6e). We show histograms of the diameters in Figure S9. Both these yields were lower than those obtained using PCP-assisted hydration. Furthermore, the GUVs obtained via electroformation were smaller than those obtained via PCP-assisted hydration, which is consistent with the observation that GUVs are small when the yields are low (Figure 2a,b).

**Figure 6.**
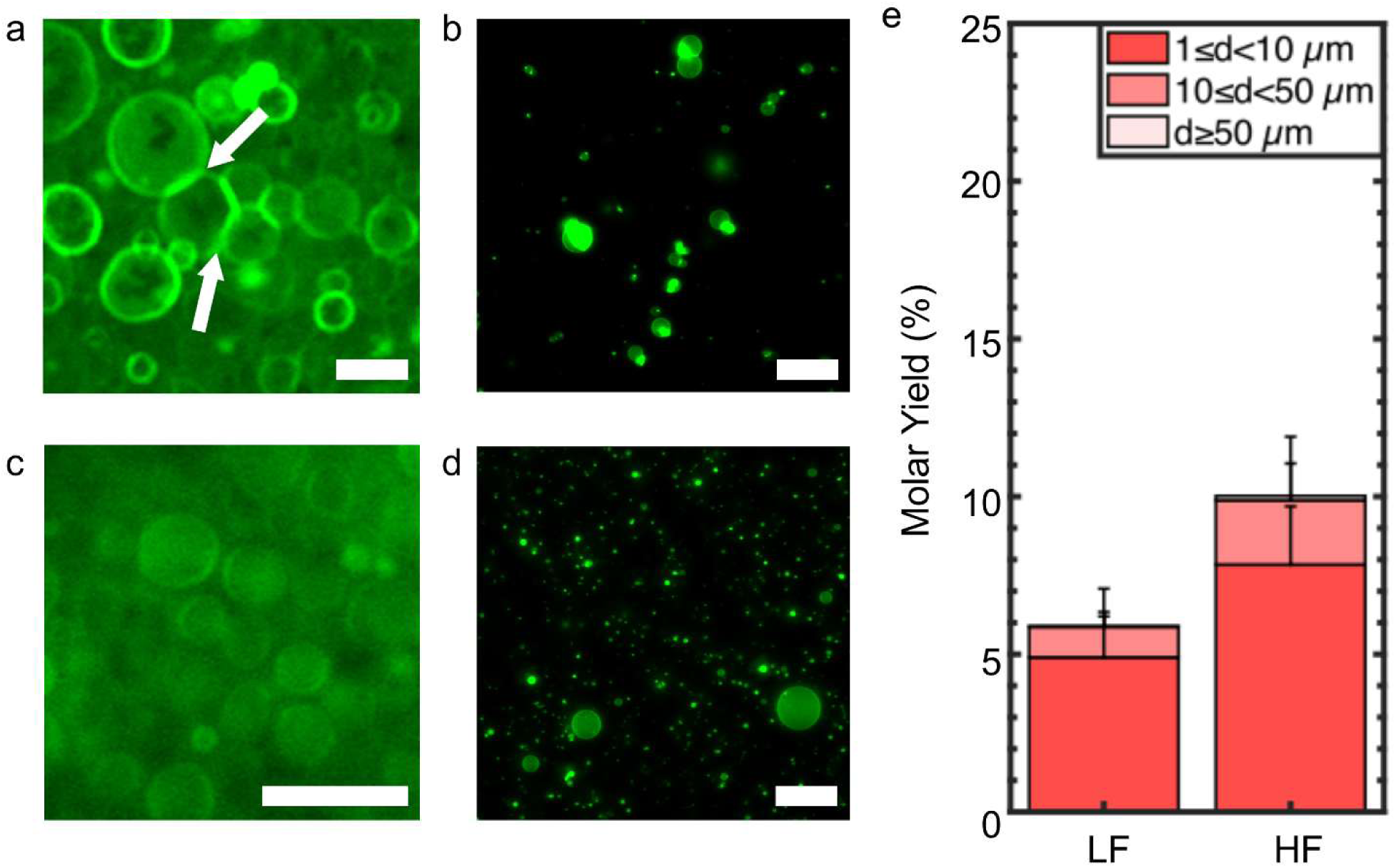
Assembly of GUVs using electroformation in salty solutions. The ITO slides were not coated with polymers. a) Image of GUV buds with 0 mol % PE2000-DSPE in the membrane on the surface of the ITO slide using the high frequency (HF) protocol. The white arrows point to regions where membranes are in adhesive contact. b) Representative single plane confocal images of the harvested objects when the membrane contained 0 mol % PEG2000-DSPE. c) Image of GUV buds with 3 mol % PE2000-DSPE in the membrane on the surface of the ITO slide using the HF protocol. d) Representative single plane confocal images of the harvested objects when the membrane contained 0 mol % PEG2000-DSPE. e) Molar yield of GUVs obtained via electroformation using low frequency (LF) or HF protocols. The stacks show the percentage of the molar yield that is comprised of GUVs with the diameters listed in the legend. The bar is an average of N=3 independent repeats. The error bars are one standard deviation from the mean. Scale bar for a,c) 20 µm. Scale bar for b,d) 50 µm.

### 2.7 The effects of lipids with repulsive headgroups on GUV assembly

We use 3 mol % of PEG2000-DSPE in our lipid mixtures to prevent aggregation of the GUV buds on the surface and the harvested GUVs in salty solutions.^[3,48]^ Some applications, such as intracellular delivery of cargo,^[49]^ may not desire the use high concentrations of PEG2000-DSPE. We thus investigate the requirements to obtain colloidally-stable isolated GUVs in salty solutions using PCP-assisted hydration. We test different mol percentages of PEG2000-DSPE and test the effects of lipids with partial molecular similarity to PEG2000-DSPE on the process of assembly. PEG2000-DSPE has both a negatively-charged deprotonated phosphate group at pH 7.4 and 2000 MW hydrophilic PEG chain. We used a lipid with a shorter PEG chain, 1,2-distearoyl-sn-glycero-3-phosphoethanolamine-N-[methoxy(polyethylene glycol)-350] (ammonium salt) (PEG350-DSPE), a lipid with a PEG chain but without a charge group, distearoyl-rac-glycerol-PEG2000 (PEG2000-DSG), and a lipid with a charge group and no PEG chain, 1,2-dioleoyl-sn-glycero-3-phospho-(1’-rac-glycerol) (sodium salt) (DOPG). We used two concentrations of DOPG, 3 mol % and 25 mol %.

In the absence of PEG2000-DSPE, similar to our results for electroformation, the buds adhere to one another on the surface (Figure 7a, white arrows). In contrast to electroformation, however, there were many more buds on the surface. GUVs that were harvested from the film without PEG2000-DSPE appear as clusters of various sizes that were composed of aggregated GUVs (Figure 7b, S10). While undesirable for biophysical applications that favor free-standing isolated GUVs, clusters of GUVs could be useful models of prototissues or cell clusters.^[50,51]^ We note that the distribution of cluster sizes can likely be tuned by controlling the surface concentration of lipids, thereby adjusting the number density of buds. Another parameter that could be tuned is the ionic strength of the solution.

**Figure 7.**
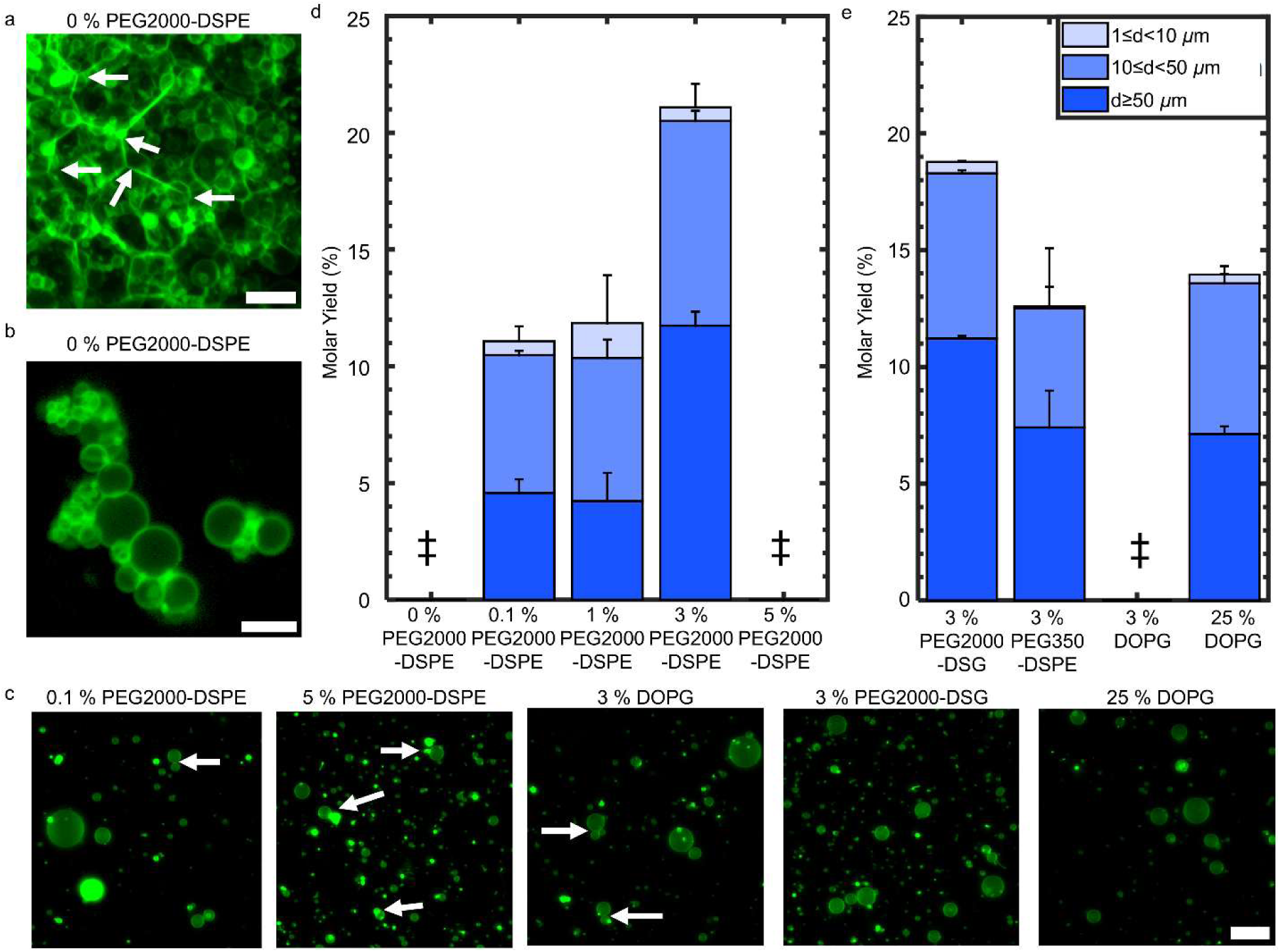
PCP-assisted hydration with membranes containing lipids with various charged and polymer-modified headgroups. PCP was prepared with ULGT agarose at the optimal polymer-NSC of 1.5 nmol/cm^2^. a) Z-projection using the “*sum slices”* method of a hydrated film with 0 mol % PEG2000-DSPE in the membrane. The white arrows point to membranes that are in adhesive contact. b) Representative single plane confocal images of the harvested objects when the membrane contained 0 mol % PEG2000-DSPE. The buds are harvested as clusters. c) Representative single plane confocal images of the harvested objects with the lipid composition and mol % listed in the labels. d) Evolution of the molar yield with PEG2000-DSPE concentration. The data for 3 mol % PEG2000-DSPE is reproduced from Figure 1 for ease of comparison. e) The molar yield of GUVs when the membrane is composed of lipids with partial molecular similarity to PEG2000-DSPE. The stacks show the percentage of the molar yield that is comprised of GUVs with the diameters listed in the legend. Each bar is the average of N=3 independent repeats. The error bars are one standard deviation from the mean. ‡ refers to conditions where yield could not be measured due to aggregation of the GUVs. Scale bar a, b) = 10 µm.

We find that even a relatively low concentration of 0.1 mol % of PEG2000-DSPE was sufficient to reduce the fraction of aggregated GUVs (Figure 7c, white arrows point to adhering GUVs, see additional images in Figure S11). Interestingly, we found that with 5 mol % PEG2000-DSPE in the membrane, the GUVs aggregated. The membrane containing 3 mol % of DOPG had aggregated GUVs while the membrane containing 25 mol % of DOPG had isolated GUVs (Figure 7c). Although low amounts of PEG2000-DSPE is sufficient to increase the fraction of isolated GUVs, we find that the yield of GUVs correlated with the concentration of PEG2000-DSPE in the membrane (Figure 7d). See Figure S12, Table S6, and Table S7 respectively for the histogram of diameters, the results of an ANOVA test, and the results of Tukey’s HSD. The yield of isolated GUVs increased with PEG2000-DPSE concentration and was highest at 3 mol %. Since the GUVs aggregated when the membranes contained 0 mol % PEG2000-DSPE or 5 mol % PEG2000-DSPE, a molar yield for the isolated GUVs could not be reliably measured. Micellar phases are favored for PEG2000-DSPE concentrations > 5 mol %.^[2,52,53]^ We speculate that micellization may decrease the stability of the GUVs to aggregation in salty solutions by reducing the effective concentration of PEG2000-DSPE in the membrane. The dependence on yield of PEG2000-DSPE was qualitatively similar to the behavior of GUVs obtained via shear-induced fragmentation in low salt solutions using gentle hydration on glass slides.^[2]^ Unlike gentle hydration, however, there is no foam-like mesophase, and the configuration of the buds on the surface did not show dramatic differences with the mol % of PEG2000-DSPE. The lipid film formed size-stratified layers of buds on the surface (Figure S13).

Increasing the concentration of PEG2000-DSPE from 0.1 to 3 mol % should decrease the magnitude of the adhesive potential between the membranes,^[2,54]^ reducing the free energy cost of budding. We propose that this result explains the steady increase in yield with PEG2000-DSPE mol %. Membranes with 3 mol % PEG2000-DSG resulted in a similar yield, 18.8 ± 0.1 %, to membranes with 3 mol % PEG2000-DSPE (Figure 7e). Membranes with 3 mol % of PEG350-DSPE resulted in a GUV yield of 12.6 ± 2.5 %. The yield with 25 mol % of DOPG was 14.0 ± 0.4 %. These two yields were statistically indistinguishable from each other and from the yield of GUVs when the membrane contained 0.1 and 1 mol % of PEG2000-DSPE. Only the compositions that had 3 mol % of PEG2000 modified lipids had yields that were significantly higher than the other compositions that we tested (all p-values < 0.01, Table S7). Taken together, these results suggest that budding in salty solutions is aided by the steric repulsion of the chain. Use of lipid modified by longer PEG chains at 3 mol % promotes more efficient budding than shorter PEG chains or other mol percents. In biomedical applications, addition of PEG2000 modified lipids to lipid formulations is routine since it confers stability against opsonization and macrophage uptake.^[55,56]^ However, in applications that do not require this stabilization, 25 mol % of DOPG appears to confer sufficient stability to the GUVs to prevent aggregation.

### 2.8 PCP-assisted hydration enables facile reconstitution of complex biochemical encapsulants

Since obtaining GUVs in salty solutions is desired to reconstitute biochemical reactions within cell-like confines and to assemble synthetic cells, we tested the ability of PCP-assisted hydration to functionally encapsulate two “difficult” biochemical reactions, cell-free transcription translation (TXTL) expression of proteins from a nucleic acid template^[14]^ and the reconstitution of actin bundles. Commercial cell-free expression platforms are proprietary mixtures of high concentrations of proteins, salts, and other promoters. Reconstitution of actin bundles requires the encapsulation of G-actin and bundling proteins in monomeric form prior to actin polymerization and bundling in GUVs.^[11–13]^

We perform PCP-assisted hydration using a commercial cell-free protein expression mixture (myTXTL^®^ Pro) with 5 nM (13.4 ng/µL) of T7 deGFP plasmid. Successful expression of the plasmid results in the formation of green fluorescent protein (GFP), which can be visualized with fluorescence microscopy. We cut the PCP into 3 mm diameter disks to scale down our typical assembly volume by 12.5× to the 12 µL volume recommended by the manufacturer. We use 0.5-mL PCR tubes for fluid management (Figure 8a). We perform PCP-assisted hydration at 4 °C. The low temperature prevents the premature expression of deGFP.^[14]^ After 1 hour, we harvested the GUV buds to obtain GUVs and then incubated the suspension at room temperature for 6 hours to allow gene expression. We then imaged the GUVs using dual-channel confocal microscopy. Figure 6b shows representative images of the harvested GUVs. The method resulted in ∼ 1 × 10^5^ GUVs in the chamber. Figure 6c shows a histogram of the luminal intensity of n = 660 GUVs (green bars). We show the results of a second independent repeat in Figure S14. The mean intensity in the lumens was 25 ± 8 arbitrary units (AU), and the lowest measured intensity was 5.5 AU. In contrast, the mean intensity of a control that was prepared and imaged identically but with the plasmid omitted was 3 ± 1 AU. The highest measured luminal intensity was 5 AU, showing that there was no overlap in intensity values between the control and deGFP expressing GUVs (Figure 8c, white bars). We conclude that 100% of the GUVs prepared with the T7 deGFP plasmid had measurable gene expression. In a second independent sample, 98% of GUVs had measurable gene expression (Figure S14). Previous reports encapsulating commercial PURE systems using the emulsion transfer technique with 10 ng/µL of template DNA resulted in 87 % of the GUVs having measurable GFP expression^[14]^, and a bead-based freeze-thaw technique with 7.4 nM template DNA resulted in 30 % of the GUVs having measurable yellow fluorescent protein (YFP) expression.^[15]^ An additional parameter that is often measured is the coefficient of variation (CV), which reflects the variation in the concentration of the expressed protein within the population of GUVs. Reported CVs of GFP intensity for the emulsion transfer technique is between 0.3 and 0.5.^[16,17]^ The CV for YFP intensity expressed in the lumens of the GUVs from the bead-based freeze-thaw technique was not reported. We obtained a CV of 0.30 for the sample shown in Figure 8 and a CV of 0.31 in Figure S14. We conclude that PCP-assisted hydration is effective at obtaining hundreds of thousands of GUVs with high expression levels and low CV while using minute quantities of the cell-free expression mixture.

**Figure 8.**
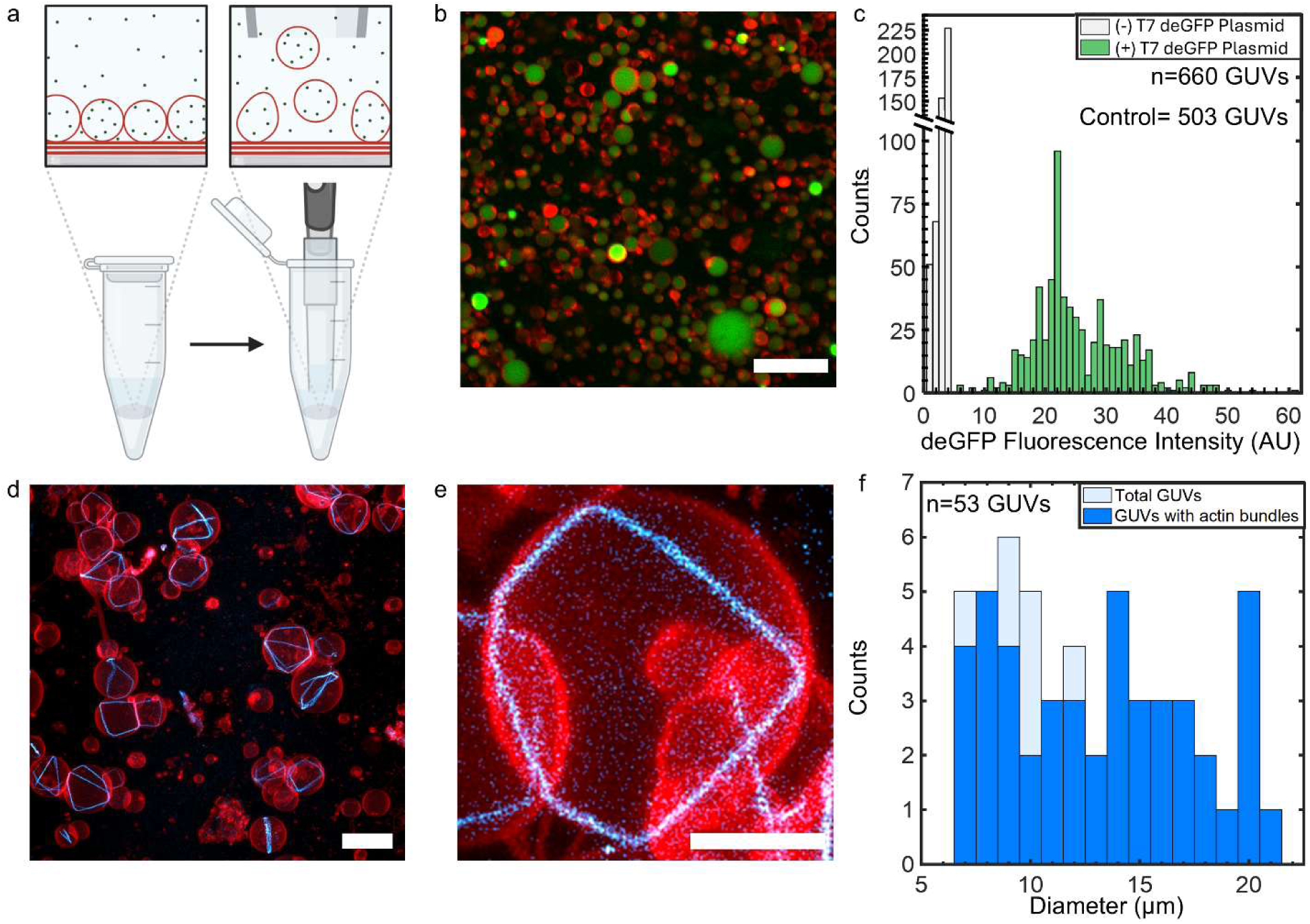
Encapsulation of complex biochemical mixtures using PCP-assisted hydration. a) Schematic of the encapsulation process of small volumes of precious mixtures in 0.5-mL PCR tubes. The zoom shows a conceptualization of the surface. The red lines are lipid membranes. The green dots represent the encapsulant. b) Representative single plane two-channel confocal image of GUVs encapsulating myTXTL cell-free expression mixture with deGFP plasmid after 6 hours of incubation. The image is an overlay of the membrane channel, false colored red, and the deGFP channel, false colored green. c) Histogram of fluorescence intensities of deGFP in the lumens of GUVs that encapsulate myTXTL cell-free expression mixture. The green bars show the distribution of luminal intensities of the expressed deGFP, n=660 GUVs. The grey bars show the distribution of luminal intensities of a negative control that lacked the deGFP plasmid, n= 503 GUVs. Two-dimensional projections of superresolution *z*-slices of GUVs with actin rings in their lumens using the “*maximum intensity projection”* algorithm. The images are an overlay of the membrane channel, false-colored red, and the actin channel, false-colored cyan. d) multiple GUVs with actin rings in their lumens. e) A zoomed-in projection of a GUV with a single actin ring. f) Stacked bar plot of the counts of GUVs with actin rings vs the diameter of GUVs. n = 53 GUVs. Scale bars a,d) 20 µm, e) 2 µm.

We next evaluated the effectiveness of the PCP-assisted hydration technique for obtaining actin bundles in GUVs. We encapsulate 12 µM actin with 1.8 µM fascin in actin polymerization buffer with the goal of forming actin rings. We use a 12 µL reaction volume and perform PCP-assisted hydration at 4 °C to reduce the premature polymerization of G-actin into F-actin during the encapsulation process.^[11]^ We acquired confocal Z-stacks and projected the three-dimensional stacks into two-dimensional images using the ‘*maximum intensity projection’* method in FIJI. We obtain GUVs that ranged in diameters from 2 µm to 20 µm. We limited our analysis to GUVs that had diameters ≥ 6.5 µm as it was difficult to discern the thin actin bundles in small GUVs. We find that 74% of GUVs had actin bundles in their lumens and that the presence of actin bundles did not depend on the diameter of the GUVs (n = 68 total GUVs counted, Figure S15). Movement of the GUVs during Z-stack acquisition, and the relatively low resolution of the 20× 1.0 NA Plan Apochromat objective that we used, made it difficult to discern the morphology of the actin bundles that were present in the GUVs. We prepared a second independent sample with 1 mol % of biotinylated lipid in the membrane and immobilized the GUVs onto the surface of a biotinylated glass coverslip using streptavidin. We then obtained superresolution images of the actin bundles that were encapsulated in the immobilized GUVs (Figure 8d-e). We find that 87% of the GUVs had actin bundles in their lumens. The presence of actin bundles did not depend on the size of the GUVs (n=53 total GUVs counted, Figure 8f). 26 % of the GUVs had single actin rings, 54 % of the GUVs had single actin rings that were not closed or had one or more branches, and 20 % of the GUVs had multiple actin rings that consisted of single actin rings or rings with branches (n=46 GUVs with actin bundles). Examples of GUVs encapsulating these structures are shown in Figure S16. This result is comparable to encapsulating actin and fascin using the transfer of emulsion droplets via the C-DICE technique, where the authors found that 12 % of the GUVs had single actin rings.^[13]^ We estimate that there were ∼ 1 × 10^4^ GUVs with actin bundles in the chamber. Furthermore, as expected, increasing the actin concentration resulted in GUVs with network-like actin bundles that could in some cases deform the membranes to form protrusions (Figure S17).^[11]^

Incorporation of biotinylated lipid, which allows attachment of biotinylated proteins^[57]^ or to surfaces using streptavidin, was straightforward. We find that it was equally easy to incorporate other functionalized lipids, such as 1,2-dioleoyl-sn-glycero-3-[(N-(5-amino-1-carboxypentyl)iminodiacetic acid)succinyl] (nickel salt) (18:1 DGS-NTA(Ni)), which have been used to attach His-tagged proteins to the membrane,^[57]^ without affecting the yields of the GUVs (Table 1).

### 2.9 Polymer-coated nanocellulose paper is stable and PCP-assisted hydration can be easily scaled up

In addition to scaling down to work with small volumes of precious samples, PCP-assisted hydration can also be scaled up to obtain large numbers of GUVs by using large pieces of PCP. We show an example of a protocol that can be performed in a typical laboratory using a 330 mm × 279 mm (13-in. × 11-in) commercial baking tray as a fluid receptacle and a piece of paper 279 mm × 178 mm (11-in × 7-in). For this experiment, we use dextran PCP since dextran has a lower cost compared to the other polymers (Table S8). We obtain 188 mL of suspension containing a total of 5.5 × 10^9^ GUVs in a single preparation (Figure 9a). The histogram of diameters is shown in Figure S18. The number of GUVs obtained is 1344× higher than a single well in a 48-well plate and 28× higher if hypothetically all 48-wells were used (Figure 9b). We create an artificial tissue-like structure with the GUV suspension. Figure 9c shows a 300 µm × 300 µm two-dimensional Z-coded projection of a tissue-like layer with a length × width × height of 5900 µm × 5900 µm × 24 µm. The tissue-like layer was obtained by pelleting the GUVs from 7.5 mL of the GUV suspension via centrifugation and then taking a 30 µL of the pellet and allowing the GUVs to sediment in an imaging chamber for 3 hours (See Experimental Section for full details). We thus could obtain ∼ 25 of these tissue-like layers from a single preparation. Notably, although the whole process took a relatively short 2.5 hours after the PCP was prepared, active user engagement was only ∼ 30 minutes during the spreading of the lipid, pouring of the hydration buffer, and the harvesting of the GUVs. The additional two hours were for evaporating traces of the solvent (1 hr) and incubating in the buffer (1 hr). Due to the short user engagement time, the method could be easily parallelized with multiple sheets of paper and baking trays.

**Figure 9.**
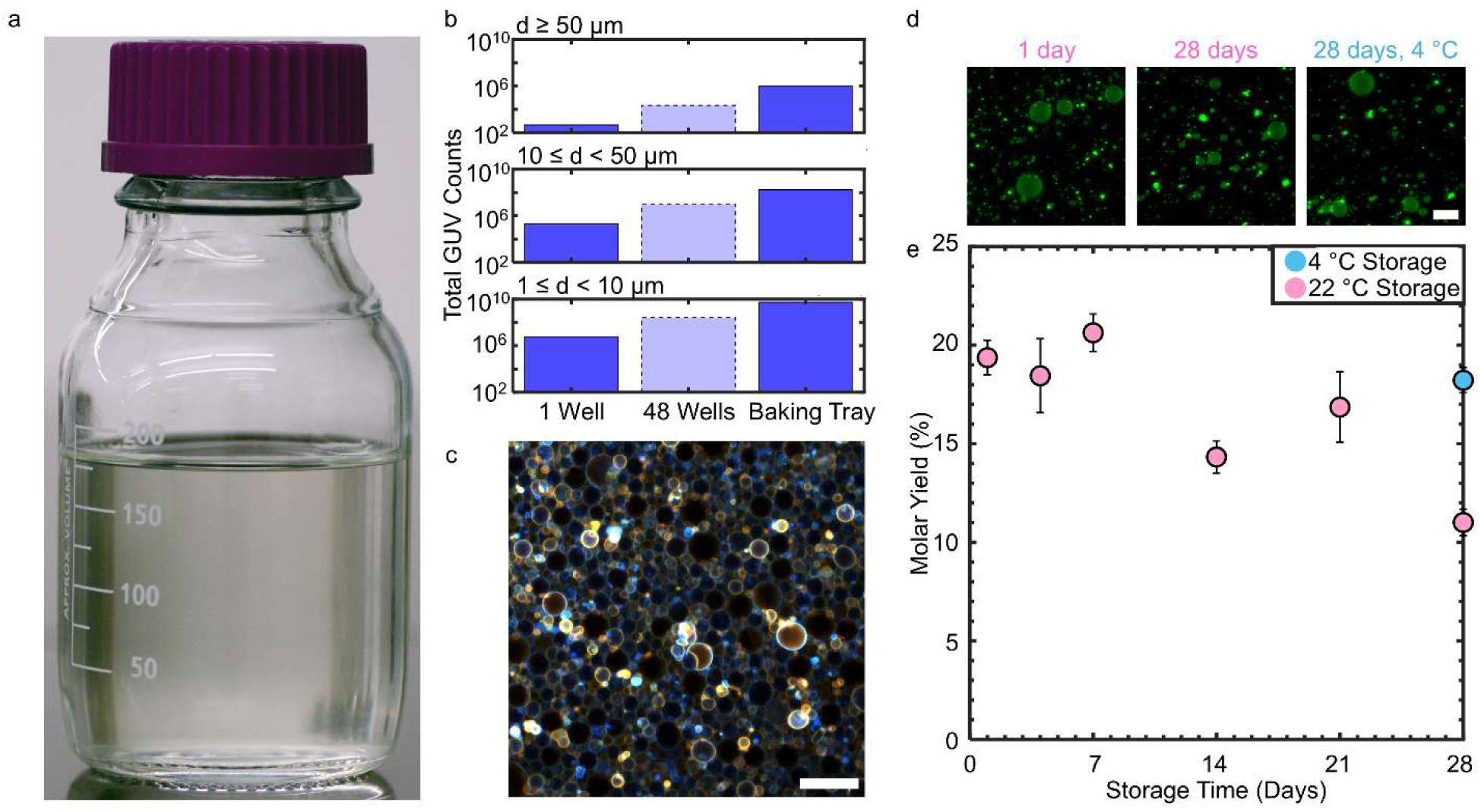
Versatility of PCP-assisted hydration. a) Photograph of a 188 mL suspension of GUVs in physiological saline obtained by performing PCP-assisted hydration using a 279 mm × 178 mm sheet of dextran PCP. The concentration of GUVs was 2.95 × 10^7^ per mL for a total of 5.54 × 10^9^ GUVs b) Bar plots of the total GUV counts from 1 well (dark blue bars with solid borders), hypothetical yield if using 48 wells (light blue bar with dashed borders), and from PCP-assisted hydration using the 279 mm × 178 mm dextran PCP (dark blue bars with solid borders). Note the logarithmic scale on the *y*-axis. Single well data is reproduced from Table 1. c) Z-coded projection of a tissue-like assembly of close-packed GUVs. d) Representative images of GUVs obtained from ULGT agarose PCP that was stored for 1 day and 28 days at 22 °C, and stored for 28 days at 4 °C. e) Plot of the molar yield of GUVs obtained using stored ULGT agarose PCP. Scale bars are 50 µm.

Since preparation of the PCP required the longest time in the protocol, ∼ 4 hours, we investigated how long a piece of ULGT agarose PCP could be used after fabrication. We find that PCP stored on a room temperature laboratory bench can be used for up to 21 days without significant loss of GUV yield or apparent reduction in the quality of GUVs that are obtained (Figure 9d-e). After 28 days of paper storage at room temperature, however, the yield falls significantly to 11.0 ± 0.7 %. The GUVs that were harvested appeared identical to those obtained at earlier times. Storing the paper at 4 °C prevented this apparent slow degradation of the properties of the PCP. There was no reduction in yield when the paper was retrieved from the refrigerator after 28 days. However, repeated removal of the PCP from the refrigerator should be avoided since it results in condensation which decreases the yield with each removal (Figure S19).

## 3 Conclusion

We demonstrate that nanocellulose paper as a substrate greatly broadens the range of polymers that support GUV assembly in physiologically relevant salt solutions, enabling the use of at least six soluble macromolecules—hyaluronic acid, dextran, carrageenan, poly-D-lysine, BSA, and DNA. By coupling the nanoscale curvature of cellulose fibers with the osmotic pressure from dissolving polymers, PCP-assisted hydration achieves robust GUV formation across a wide temperature range (4–45 °C). The method also offers practical advantages: large PCP sheets can be prepared within hours for bulk GUV assembly in hundreds of milliliters of buffer or adapted for small-scale formats such as well plates and PCR tubes. Prepared PCP remains stable for at least 21 days, enabling flexible experimental workflows. We anticipate that PCP-assisted hydration, and the diverse polymer chemistries it accommodates, will become a versatile platform for GUV assembly under physiological conditions, advancing applications in biophysics, synthetic biology, and biomedical engineering.

## 4 Experimental Section Materials

We purchased premium plain glass microscope slides (75.2 mm × 25.4 mm, Fisherbrand, Catalog number: 12-544-1), full-frosted microscope slides (Fisherbrand, Catalog number: 12-550-401) 48-well polystyrene tissue culture plates (Fisherbrand, Catalog number: FB012930), Fisherbrand™ Ultra-Clean™ Supreme Aluminum Foil, glass coverslips (Corning, 22 mm × 22 mm), Thermo Scientific Nitrocellulose Membranes, 0.45 μm (Catalog number: 88025), Falcon™ Bacteriological Petri Dishes with Lid, 2 mL disposable glass Pasteur pipettes, dropper bulb, 4 mL glass vials, 50 mL Falcon centrifuge tubes, and Eisco™ Mixed Set of Glass Delivery Tubes from Thermo Fisher Scientific (Waltham, MA). We purchased indium tin oxide (ITO) coated-glass slides (25 × 25 mm squares, surface resistivity of 8−12 Ω/sq) from Sigma-Aldrich (St. Louis, MO). We purchased Grafix shrink film (8.5-in × 11-in) and a 330 mm × 279 mm (13-in. × 11-in.) baking tray (Wilton) from Amazon.com (Seattle, WA, USA). We purchased acid-free artist grade tracing paper (Jack Richeson Tracing Pad Paper 9 x 12 50 Sheets) from an online vendor Brushes and More.

### Chemicals

We purchased sucrose (BioXtra grade, purity ≥ 99.5%), glucose (BioXtra grade, purity ≥ 99.5%), casein from bovine milk (BioReagent grade), agarose type IX-A: ultra-low gelling temperature (Catalog number: A2576, molecular biology grade), agarose: low gelling point (Catalog number: A9414, molecular biology grade), deoxyribonucleic acid, single stranded from salmon testes (Catalog number: D7656), carrageenan (Catalog number: C1013), dextran from *Leuconostoc spp*. (Catalog number: 09184, MW: 100,000), bovine serum albumin (Catalog number: A3059), hyaluronic acid sodium salt from *Streptococcus equi* (Catalog number: 40583, MW: 8,000 _–_ 15,000) and (Catalog number: 96144 MW: 70,000 – 120,000), poly(vinyl alcohol) (Catalog number: 363138, MW: 31,000-50,000, 96-99% hydrolyzed) and (Catalog number: 363065, MW: 146,000-186,000 99+% hydrolyzed), streptavidin from *Streptomyces avidinii*, and DTT (OmniPur^®^ Grade ≥ 99.4%) from Sigma-Aldrich (St. Louis, MO). We purchased dextran from *Leuconostoc mesenteroides* (Catalog number: J62775, MW: 6,000) from Alfa Aesar (Ward Hill, MA). We purchased chloroform (ACS grade, purity ≥ 99.8%, with 0.75% ethanol as preservative), 96% ethanol (molecular biology grade), 200 proof ethanol (molecular biology grade), acetone (ACS grade), and Invitrogen 10× phosphate buffered saline (PBS) (pH 7.4, 0.2 µm filtered, 1.37 M sodium chloride, 0.027 M potassium chloride, 0.080 sodium phosphate dibasic, 0.020 M potassium phosphate monobasic) from Thermo Fisher Scientific (Waltham, MA). We obtained 18.2 MΩ Type I ultrapure water from a Milli-Q® IQ 7000 Ultrapure Lab Water System (Burlington, MA). We purchased 1,2-dioleoyl-*sn*-glycero-3-phosphocholine (18:1 (Δ9-cis) PC (DOPC)), 23-(dipyrrometheneboron difluoride)-24-norcholesterol (TopFluor-Chol), cholesterol (ovine wool, >98%), 1,2-dipalmitoyl-*sn*-glycero-3-phosphocholine (16:0 PC (DPPC)), 1,2-dioleoyl-*sn*-glycero-3-phosphoethanolamine-N-(lissamine rhodamine B sulfonyl) (ammonium salt) (Rhod-DOPE), 1,2-distearoyl-*sn*-glycero-3-phosphoethanolamine-N-[biotinyl(polyethylene glycol)-2000] (ammonium salt) (PEG2000-Biotin-DSPE), 1,2-dioleoyl-sn-glycero-3-[(N-(5-amino-1-carboxypentyl)iminodiacetic acid)succinyl] (nickel salt) (DGS-NTA(Ni)), distearoyl-rac-glycerol-PEG2000 (PEG2000-DSG), 1,2-dioleoyl-sn-glycero-3-phospho-(1’-rac-glycerol) (sodium salt) (DOPG), 1,2-distearoyl-sn-glycero-3-phosphoethanolamine-N-[methoxy(polyethylene glycol)-350] (ammonium salt) (PEG350-DSPE), and 1,2-distearoyl-*sn*-glycero-3-phosphoethanolamine-N-[methoxy(polyethylene glycol)-2000](ammonium salt) (PEG2000-DSPE) from Avanti Polar Lipids, Inc. (Alabaster, AL). We purchased a myTXTL Pro Cell-Free Expression Kit from Daicel Arbor Biosciences (Ann Arbor, MI). We purchased actin (> 99% pure, rabbit skeletal muscle), actin: HiLyte™ Fluor 488 Labelled (>99 % pure, rabbit skeletal muscle) fascin-1 (wild-type, human-recombinant), actin polymerization buffer (10× stock), and general actin buffer (1× stock) from Cytoskeleton, Inc. (Denver, Co).

### Procedure for cleaning glass slides, ITO-coated slides, and nanocellulose paper

We clean eight 25 mm × 75 mm glass slides by sonicating in a Coplin staining jar sequentially for 10 minutes each in acetone, 200 proof ethanol, and Type I ultrapure water.^[2]^ We dry the slides under a stream of nitrogen and place the slides in a 65 °C oven for 2 hours.

We cut a single 483 mm × 610 mm (19 in × 24 in) sheet of nanocellulose paper into 1/8^th^ sections and place the sections in 250 mL of 96 % molecular biology grade ethanol in a glass crystallizing dish (45 mL, dish diameter × height 40 mm × 25 mm) for 15 minutes. The liquid in the dish was manually agitated occasionally. After 15 minutes the ethanol was discarded and the process repeated once. 250 mL of Type I ultrapure water was then added to the dish, swirled three times, and then discarded. This process was repeated an additional three times. Then 250 mL of Type I ultrapure water was added to the dish and allowed to soak for a further 15 minutes with occasional manual agitation. The water was then discarded, and the process was repeated once. Then the individual paper sections were removed and placed on an aluminum foil. We then placed another piece of aluminum foil loosely on top of the foil to prevent dust particles from settling on the wet paper sheets. We placed the foil with wet paper in a 65 °C oven overnight. The next day we remove the paper and store it in a closed 150 mm diameter polystyrene Petri dish on a lab bench until further use.

### Preparation of polymer solutions

The concentration of polymer solutions, the temperature, the method of agitation, and the approximate time for dissolution are shown in Table S9. We weigh the mass of polymer using an analytical balance. We then dissolve the polymer in Type I ultrapure water in a 2 mL PCR tube or a 15 mL conical centrifuge tube depending on the volume of polymer solution we needed. We prepare 1 mL (hyaluronic acid, poly-D-lysine, salmon DNA, BSA) or 10 mL (agarose, PVA, dextran, carrageenan) of polymer solution. Agarose, PVA, and carrageenan were heated using a dry heating block (Thermo Scientific Multi-Blok Heater) set at the desired temperature to aid dissolution (Table S9). For experiments to determine the optimal polymer-NSC, the polymer solutions were diluted with isotemperature Type I ultrapure water. For the fabrication of large pieces PCP, the polymer solutions were prepared at the concentration needed to obtain the desired polymer-NSC, for example, 0.3 % w/w for ULGT agarose and 4.85 % w/w for dextran. Polymer solutions were applied immediately at the preparation temperature onto paper preheated to 40 ° C. We prepare fresh polymer solutions each time we prepare paper.

### Fabrication of PCP and PCG substrates

We place the cleaned glass slides on a Parafilm piece on a hotplate set at 40 °C and deposit the polymer solution using a micropipette (e.g. 1,162 µL of the ULGT agarose solution at various % w/w). We spread the solution evenly with the side of a 1000 µL pipette tip. The glass slides were left on the hotplate for 4 hours to dry. The PCG substrates were stored face up in a clean Petri dish and used within 24 hours. Polymer coated frosted glass slides were fabricated using the same method.

We prepare PCPs of different sizes. We cut the cleaned nanocellulose paper to 22 mm × 22 mm squares when using relatively expensive polymers or when we planned to do a limited number of experiments. We place the pieces of paper on a piece of Parafilm on a hotplate set at 40 °C. We then apply 300 µL of the polymers at various % w/w on the substrates. The polymer solution was evenly spread on the substrates with the side of a 1000 µL pipette tip. Polymer coated nitrocellulose membranes were fabricated using the same method. We coat a whole strip of the cleaned 76 mm × 114 mm paper when we perform multiple experiments and when we store the paper. We apply 5.4 mL of warm 0.3 % w/w ULGT agarose solution using a 1000 µL micropipette in a serpentine pattern across the length of the paper. Then, the 60 mm section of an L-shaped glass delivery tube (180 mm × 60 mm) was passed four times back and forth along the 114 mm length of the paper to spread the solution evenly, careful not to allow any polymer solution to spill over the sides of the paper. The substrates were left on the hotplate for 4 hours. PCP was stored face up in a clean Petri dish.

Commonly available lab hot plates could not accommodate the 279 mm × 178 mm (11-in. × 7-in). piece of nanocellulose paper for the experiments with the baking tray. We thus place a sheet of aluminum foil as a backing on a baking tray followed by the nanocellulose paper. To obtain a polymer-NSC of 4.3 nmol/cm^2^, we deposit a total of 4.25 mL of a 4.85 % w/w solution of dextran onto the paper. Using a bulb and glass Pasteur pipette, we deposited 3 mL of the solution onto the paper in a serpentine pattern on the left 2/3^rd^ of the paper along the 178 mm width. The 180 mm handle of the L-shaped glass delivery tube was moved up and down once in a smooth controlled motion to spread the dextran solution evenly. We next apply 1.25 mL of the dextran solution onto the right 1/3^rd^ of the paper in a straight line. We complete the coating process by moving the 60 mm side of the glass delivery tube up and down once along the 178 mm width. We then placed the baking tray with the wet paper in an oven set to 40 °C to dry for 4 hours.

### Buffers

All hydration and sedimentation buffers are made as needed from stock solutions of 1 M sucrose, 1 M glucose, and 10 × PBS and used within 1 hour. 1 M sucrose and glucose stock solutions were stored in a 4 °C fridge and used for a maximum of two weeks. If solutions are used past this time, we find that the yield of the controls had a high likelihood of being low. Occasionally, even when all protocols are followed the yield falls. In these cases, remaking the buffers and performing the controls restore yields. We use 150 µL of 1× PBS + 100 mM sucrose as the hydration buffer for PCP-assisted hydration performed at 4 °C, 22 °C, 37 °C for 1 hour and at 45 °C performed for 20 minutes. We use 160 µL 0.97× PBS + 97 mM sucrose as the hydration buffer for PCP-assisted hydration performed at 45 °C for 1 hour to account for the 10 µL of buffer that evaporates. All buffers were warmed to the desired temperature for 1 hour using a dry heating block (Multi-Blok Heater, Thermo Scientific) or chilled to the desired temperature in an Isotemp™ low temperature incubator (model number: 3724, Fisher Scientific, Waltham, MA) before use. For reconstitution of actin we use 1× general actin buffer (5 mM Tris-HCl pH 8.0 and 0.2 mM CaCl_2_). Actin polymerization was triggered by adding 1/10^th^ of 10× actin polymerization buffer (500 mM KCl, 20 mM MgCl_2,_ 50 mM guanidine carbonate pH 7.5, and 10 mM ATP in 100 mM Tris, pH 7.5) to obtain a final concentration of 1× (50 mM KCl, 2 mM MgCl_2,_ 5 mM guanidine carbonate pH 7.5, and 1 mM ATP in 10 mM Tris, pH 7.5).

### Deposition of lipids

We deposit lipids as described previously.^[1,3]^ We cut circular disks of the PCP using a circle hole punch (EK Tools Circle Punch, 3/8 in.). We deposited 10 µL of the lipid solution onto the polymer-coated side of the PCP evenly using a glass syringe (Hamilton) while holding the paper with clean metal forceps. The lipid-coated PCP disks were placed on a clean glass slide and into a standard laboratory vacuum desiccator for 1 hour to remove traces of organic solvent prior to hydration.

### Procedure for PCP-assisted hydration and PCG-assisted hydration

We perform PCG-assisted hydration as previously described.^[2]^ We perform PCP-assisted hydration as previously described for the PAPYRUS technique with minor modifications.^[1,3]^ 48-well plates were warmed on a hotplate or chilled in a low temperature incubator for 1 hour if hydration was not performed at room temperature. Then, the lipid-coated PCP was placed into the wells of the 48-well plate and allowed to equilibrate for 5 minutes. Next, we add the isotemperature hydration buffer to the wells and allow the samples to hydrate for 1 hour. For experiments probing the effect of incubation time, we changed the incubation time to 20 mins, 2 hours, or 24 hours.

### Procedure for harvesting the GUVs

We harvest GUVs from the substrates as previously described at the temperature of assembly.^[2]^ We pipette 100 µL of the hydrating solution with a cut 1000 µL pipet tip on 6 different regions of the PCP to cover the whole area. On the 7^th^ time, we aspirate all the GUV containing solution and transfer the liquid into a 0.5-mL PCR tube. Any GUV solutions above 22 °C are allowed to cool in a PCR tube rack at 22 °C before imaging.

### Control procedures

We normalize the concentration of the lipid solution that we prepare to control batch-to-batch variations of the mass of DOPC powder obtained from the manufacturer. We prepare GUVs using 10 µL of a nominal 1.0 mg/mL of DOPC:TopFluor-Chol 99.5:0.5 using the standardized PAPYRUS procedure^[1,4]^ using freshly prepared buffers and clean nanocellulose paper. We measure the molar yield and the distribution of diameters. If the yield was below 33 ± 2 %, we infer that the mass of DOPC was different from the stated nominal mass. To correct this difference, we perform the PAPYRUS technique by applying 10 µL of the lipid solution at a nominal concentration of 0.8 mg/mL, 1.0 mg/mL, and 1.2 mg/mL. We calculate the molar yield assuming an applied lipid amount of 10 µg. We renumber the concentration of the lipid solution that resulted in a yield closest to a range of 32 ± 2 % as 1 mg/mL. We find that approximately 1 in 10 DOPC lipid vials from the manufacturer had an apparent lipid mass that was sufficiently different to require this correction.

To minimize technique-dependent variability, we ensure that the experimentalist can perform the PAPYRUS technique three independent times on three separate days and obtain a molar yield of 33 ± 2 % and a distribution of sizes similar to those previously published. Reproducing previously published results serves as a check for the experimentalist’s skill and care before trying new conditions with unknown yields. Lower than expected yields with validated lipid and buffers were often due to incorrect lipid deposition technique (allowing the lipid to coat the underside of the paper, for example) or poor harvesting technique. Good harvesting technique results in almost all GUV buds to be harvested from the surface (Figure S2).

### Electroformation procedure

We assembled GUVs from DOPC + 3 mol % PEG2000-DSPE through electroformation on ITO-coated glass. We use the same lipid deposition procedure to be consistent with the PCP and PCG samples. We affixed circular PDMS gaskets (inner diameter × height = 12 × 1 mm) around the dried lipid film to construct a barrier for hydration. We add 150 mM of 1× PBS + 100 mM sucrose and seal the chamber using a second ITO-coated glass slide. The ITO surfaces were then connected to the leads of a function generator (33120A Agilent) using conductive copper tape. We applied a sinusoidal AC field at a field strength of 1.5 V/mm peak-to-peak at a frequency of 10 Hz for 2 hours. For a second sample, we used a modified frequency protocol optimized for high salt conditions. We applied a sinusoidal AC field at a frequency of 500 Hz while varying the field strength. We use a field strength of 0.106 V/mm peak-to-peak for 5 minutes, 0.940 V/mm peak-to-peak for 20 minutes, and 2.61 V/mm for 90 minutes consecutively.

### Imaging GUVs to obtain yields

Imagining of the harvested free-floating GUVs was conducted as previously described.^[4]^ We construct imaging chambers by placing poly(dimethylsiloxane) (PDMS) chambers with dimensions of 5.9 mm × 5.9 mm × 1 mm (length × width × height) on glass microscope slides. We passivate the chamber with 1 mg/mL casein in PBS to prevent the rupture of GUVs on the surface of bare glass. We thoroughly rinse the chambers with ultrapure water after passivation by gently pouring 50 mL of water from a 50 mL Falcon tube over the chambers three times. We measure the osmolarity of a 20 µL aliquot of the GUV suspension and a 20 µL aliquot of our standard sedimentation buffer, 1×PBS + 100 mM glucose, using a freezing point depression multi-sample osmometer (Model 2020, Advanced Instruments, USA). The standard sedimentation buffer served as an internal control. The osmolarity of the GUV suspension varied slightly from sample-to-sample due to the dissolution of the polymer. Osmotic pressure of the GUV solution can range from 368 mOsm/kg H_2_O to 438 mOsm/kg H_2_O. The osmolarity of the standard sedimentation buffer was 375 mOsm/kg H_2_O. We adjust the concentration of the glucose to ensure that the sedimentation buffer was isoosmolar with the GUV suspension. We fill the passivated chamber with 58 µL of the isoosmolar sedimentation buffer. We pipette 2 µL aliquot of the harvested GUV suspension in the middle of the chamber, aspirate 2 µL of the solution and deposit in the middle of four quadrants and then the mid-section of the chamber to ensure even distribution of the GUVs. We cover the sedimentation chambers with a glass coverslip and allow the GUVs to sediment for 3 hours in a covered 150 mm Petri dish with a moistened Kimwipe to minimize evaporation. We use an upright confocal laser-scanning microscope (LSM 880, Axio Imager.Z2m, Zeiss, Germany) with a 10× Plan-Apochromat objective with a numerical aperture of 0.45 to capture images. We image using an automated tile scan and autofocus routine (49 images, 5951.35 µm × 5951.35 µm, (3212 pixels × 3212 pixels) to capture the entire area of the chamber. For vesicles labeled with TopFluor-Chol, we use a 488 nm argon laser. We set the pinhole to 15.16 Airy units which gave a confocal slice thickness of 79.3 µm. For phase-separated vesicles we capture dual-channel images. We use a 488 nm argon laser and a 561 nm diode-pumped solid-state laser. We image using the line-switching scanning mode. We set the pinhole of the rhodamine channel to be 14.57 Airy units which gives a confocal slice thickness of 79.4 µm. We set the pinhole of the TopFluor-Chol channel to be 17.76 Airy units which gives a confocal slice thickness of 79.4 µm. We normalize the laser power by measuring the intensity of 10 vesicles with diameters between 9 and 11 µm from 3 images using the line tool in FIJI. We adjust the laser power so that the intensity was within a range of 59 ± 3 arbitrary intensity units.

### Image Processing and Calculations of the Distribution of Sizes and Yields

We processed and analyzed the images using a custom MATLAB routine as previously described.^[1–4]^ The routine selects GUVs from thresholded objects based on the coefficient of variation (CV) of the intensity values. Objects that fell outside of 1.75 times the full width at half the maximum (FWHM) of the highest peak in the histogram of CV values were classified as not GUVs. Additionally, objects that had more than 10% of their pixel values higher than 200 on a 256 intensity units scale were classified as not GUVs. Counts of GUVs were normalized per μg of lipid deposited on the substrate. We calculate the molar yield, expressed as a percentage, using 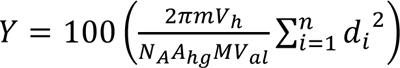. In this equation, *m* is the molecular weight of the lipid, *V*_h_ is the volume of the harvested GUV suspension, *N*_A_ is Avogadro’s number, *A*_hg_ is the headgroup area of the lipid, *M* is the mass of lipid deposited on the surface, *V*_al_ is the volume of the aliquot in the imaging chamber, *n* is the number of GUVs in the imaging chamber, and *d_i_* is the diameter of vesicle *i*.

### Quantification of the phase fraction

The domains in the vast majority of the GUVs had coarsened to form a single L_d_ and single L_o_ domain. The GUVs sediment with random orientations to the bottom of the chamber. We count GUVs with domains oriented at a 90° angle to the imaging plane (see Figure S20 for representative examples) from two images. We measure the diameter of the GUV, *d*, and the height of the spherical cap, *h* that compose the L_d_ phase in the L_d_ channel using FIJI.^[58]^ The fraction of the L_d_ phase relative to the GUV was 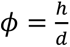. We report the value as an average ± 1 standard deviation in the main text.

### Statistical Analysis

All statistical analyses were performed using MATLAB. All experiments were repeated three independent times unless otherwise stated. For the experiments shown in Figure 7, we conducted a balanced, one-way analysis of variance (ANOVA) to determine the statistical significance of the mean yields. We then conduct a posthoc Tukey’s honestly significant difference (HSD) to determine the statistical significance between groups. When the means of two samples are compared, we conduct a Student’s *t*-test.

### Determination of optimal polymer-NSC

To determine the optimal polymer-NSC we image the configuration of the film after 1 hour of hydration. We typically prepare 6 samples by depositing 300 µL of 0.1, 0.3, 0.5, 0.7 and 1.0 % w/w of polymer on a 22 × 22 mm square of paper. We use an upright confocal laser-scanning microscope (LSM 700 Axio Imager.Z2m, Zeiss, Germany) with a 10× Plan-Apochromat objective with a numerical aperture of 0.45 to examine and capture images. In general, surfaces with polymer-NSCs that have only lipid bilayer stacks and with minimal buds will not have high yields. These polymer-NSCs are not considered further. For surfaces that have apparent buds, the buds are examined for evidence of dewetting patterns to distinguish GUV buds from pseudobuds. Surfaces that show low amount of pseudobuds and high number of GUV buds are chosen as having the optimal polymer-NSC. On some surfaces, determining which of the two surfaces with adjacent polymer-NSCs had a higher number of buds instead of pseudobuds was challenging. For these surfaces, the buds are harvested, the molar yield of GUVs is measured, and the higher yielding surface is considered to have the optimal polymer-NSC. Only the yield at the optimal polymer-NSC is reported in Table 1. Additional yields that we measured are provided in Table S2-S5, for completeness.

For the images in Figure 4 we use a 20× Plan-Apochromat objective with a numerical aperture of 1.0. We capture a Z-stack (135 images, 151.8 µm × 151.8 µm, 1272 pixels × 1272 pixels, slice thickness 1.5 µm, slice interval 0.745 µm) of a region that was representative of the surface and image from the surface of the substrate to 99.87 µm above the surface. We used the ‘*reslice*’ algorithm to obtain *x*–*z* slices from the confocal Z-stacks. The contrast of the Z-stacks is enhanced to show dim features and bright features by using the ‘*enhance contrast > equalize histogram’* option in FIJI. Non-contrast enhanced images are shown in the supporting information. We obtain *x*–*y z*-projections by using the ‘maximum intensity projection’ algorithm of the first 18.62 μm or by using the ‘sum slices’ algorithm of the first 99.87 µm.

### myTXTL expression of deGFP

We deposit 10 µg of lipid on a 9.5 mm diameter disc of PCP. We place the lipid coated PCP in a standard laboratory desiccator to remove traces of organic solvent. We use a circular hole punch (AIEX, 3mm diameter), to punch out two 3 mm disks. We punch the circular disk with the lipid coated side facing down ensuring that the lipid does not transfer to the punch. We use one 3 mm disk of lipid-coated PCP per PCR tube for a total of two samples. We prechill the 0.5-mL PCR tube in a 4 °C incubator for 1 hour before placing the lipid-coated PCP face up into the tube. We allow the paper to incubate for a further 5 minutes in the 4 °C incubator. Working on ice, we mix 12 µL of the myTXTL mix as described in the myTXTL manual.^[59]^ We place the mixture on top of the paper, close the tube, and incubate for 1 hour at 4 °C to prevent premature expression. While still working in the incubator, we harvest by pipetting 10 µL of the hydrating solution with a cut 100 µL pipet tip 6× on the PCP to cover the whole area. We remove the tubes from the incubator and pool the two samples together in one 2 mL Eppendorf tube. We allow the tube to sit for 5 minutes to come to room temperature and add 80 µL of room temperature 2.8× PBS for a total volume of ∼100 µL. This concentration of PBS was isomolar to the myTXTL mixture (∼845 mOsm/kg) which ensured osmotic balance of the GUVs. We centrifuge the pooled samples for 60 minutes at 100×g in a swing bucket clinical centrifuge (AccuSpin 8c) equipped with 3D printed adapters for 2 mL Eppendorf tubes. We then remove 80 µL of the supernatant and resuspend the pelleted GUVs in the remaining solution by pipetting 6 times. We place the suspension in a 3.5 mm circular PDMS imaging chamber, place a coverslip, and incubate at room temperature (22 °C) for 6 hours. For the negative control, we followed the same protocol but omitted the T7 deGFP control plasmid from the reaction mixture.

### myTXTL expression imaging and image processing

We obtain confocal images of the GUVs using an upright confocal laser-scanning microscope (LSM 880, Axio Imager.Z2m, Zeiss, Germany) with a 20× Plan-Apochromat objective with a numerical aperture of 1.0. We capture images with an area of 212.55 µm × 212.55 µm, (1784 pixels × 1784 pixels) using a pinhole of 1.51 A.U. for the 488 nm channel (collecting emissions from 494 nm to 555 nm) and 1.24 A.U. for the 561 nm channel (collecting emissions from 570 nm to 709 nm). We measured the fluorescence intensity of the lumen of all the GUVs from 1 image per sample using FIJI. We draw circular ROIs wholly within the dark lumen of the GUVs and avoid selecting the bright pixels that correspond to the membrane. We ignore out of focus GUVs and bright objects which are not GUVs. For the negative control, we measured the intensity of GUVs in 5 images to get ∼500 GUVs. We estimate the total number of GUVs in the chamber by dividing the number of GUVs in an image and multiplying with the total area of the chamber.

### Actin encapsulation

We deposit 10 µg of DOPC:PEG2000-DSPE:PEG2000-Biotin-DSPE:Rhod-DOPE 95.9:3:1:0.1 mol %. on a 9.5 mm diameter disc of PCP. We place the lipid-coated PCP in a standard laboratory desiccator to remove traces of organic solvent. We use a circular hole punch (AIEX, 3 mm diameter), to punch out three 3 mm disks of lipid-coated PCP. We punch the circular disk with the lipid-coated side facing down ensuring that the lipid does not transfer to the punch. We use one 3 mm disk per PCR tube for a total of three samples. To minimize temperature gradients that can hasten the polymerization of actin, we then work in a 4°C cold room for the duration of the hydration step. We place the PCR tubes containing the lipid coated-PCP on a laboratory bench in the cold room for at least 5 minutes to allow them to reach 4 °C. Working on ice, we prepare actin mix consisting of 90:10 actin: HiLyte™ Fluor 488 actin, fascin, DTT, and sucrose in general actin buffer. We then add the actin polymerization buffer and pipette 6 times to mix thoroughly. We quickly aspirate the well-mixed mixture and add 12 µL to each lipid-coated PCP disk in the PCR tubes. The final concentration of the mix shown in Figure 3 is 12 µM actin, 1.8 µM fascin, 1 mM DTT, 300 mM sucrose, 1× general actin buffer and 1× polymerization buffer. We wrap the PCR tubes with aluminum foil and allow the samples to incubate for 1 hour at 4 °C. We harvest by pipetting 10 µL of the hydrating solution with a cut 100 µL pipet tip 6 times on the PCP to cover the whole area. We pool the three samples together in one 2 mL Eppendorf tube and add 200 µL of isomolar sedimentation buffer made of 1× general actin buffer, 1× actin polymerization buffer, 1 mM DTT and 300 mM glucose for a total volume of ∼230 µL. We centrifuge the pooled samples for 20 minutes at 100×g in a swing bucket clinical centrifuge (AccuSpin 8c) equipped with 3D printed adapters for 2 mL Eppendorf tubes. We then remove 210 µL of the supernatant and resuspend the pelleted GUVs in the remaining 20 µL of buffer by pipetting gently 6 times.

### Actin imaging and image processing

GUVs imaged on the Zeiss LSM880 were not bound to the glass coverslip. We add 20 µL of the GUV suspension into a circular 3.5 mm diameter PDMS chamber and cover the chamber with a coverslip. We imaged the samples at 3, 4, and 24 hours and found no difference in actin behavior. We found locations with multiple GUVs and image confocal Z-stacks with a 20× Plan-Apochromat objective with a numerical aperture of 1.0 starting from the surface of the glass slide to 2 slices above the GUV that was furthest away from the surface. We use a pinhole of 1.01 A.U. for the 488 nm channel (slice thickness 1.33 µm, slice interval 0.667 µm, collecting emissions from 494 nm to 555 nm) and 0.83 A.U and for the 561 nm channel (slice thickness 1.4 µm, slice interval 0.667 µm, collecting emissions from 570 nm to 709 nm). The image size was 592 × 592 pixels. GUVs imaged on the Nikon Ti2-E were bound to the coverslip through biotin-streptavidin linkers. For bound GUVs, we prepare biotin functionalized coverslips as previously described.^[48]^ We place 3.5 mm circular PDMS chambers on the functionalized coverslips and allow 10 µL of 0.1 mg/mL streptavidin to incubate in the chamber for 15 minutes. We then wash the chamber 6 times with 1× PBS, then 2 times with the sedimentation buffer. We add 20 µL of the GUV suspension into the chamber and allow the GUVs to sediment for 3 hours in a closed 150 mm Petri dish with a reservoir of water to prevent evaporation. We then remove unbound GUVs by removing 10 µL of the solution from the top of the sedimentation chamber, careful not to contact the surface of the coverslip. We then add 10 µL of sedimentation buffer and seal the chamber with a glass slide. We imaged the sample after 3 hours. We use an inverted microscope (Nikon Ti2-E with AxR point scanning confocal) equipped with an NSPARC detector, a CFI Plan Apo Lambda S 60XC Silicone Immersion Objective with a numerical aperture of 1.3 to capture confocal Z-stacks. We capture confocal Z-stacks from the surface to 2 slices above the highest GUV using a pinhole of 2 A.U. The images were 2048 pixels × 2048 pixels with a slice interval 0.316 µm.

Image processing was performed using FIJI. We analyzed GUVs > 6.5 µm. GUVs were binned based on the diameter with GUVs edges of 0.5 µm below and 0.5 µm above the nearest whole number. We identified bundles in the GUV lumens manually. When the GUVs were not bound to the coverslip, it was difficult to discern whether the actin bundles were forming single actin rings, single actin rings with breaks or branches, or multiple actin rings. Therefore, we do not separate the actin bundle conformation into categories for this sample. For the superresolution images, GUVs with a single actin bundle that appeared to completely encircle the membrane of the GUV were classified as “single actin rings”, GUVs with a single actin bundle that encircled the membrane of the GUV but had gaps or with a single actin bundle that encircled the membrane of the GUV but had one or more branches were classified as “single actin rings that were not closed or had one or more branches” and GUVs with multiple actin bundles that encircled the membrane of the GUVs were classified as “multiple actin rings”. We estimate the total number of GUVs in the chamber by dividing the number of GUVs in an image and multiplying with the total area of the chamber. The images shown in Figure 8 are two-dimensional projections of three-dimensional confocal Z-stacks using the ‘*maximum intensity projection’* algorithm in FIJI.

### Large scale-assembly of GUVs using PCP-assisted hydration

We adapt the procedure for reported in reference ^[4]^ with modifications. The lipid solution consisted of DOPC: PEG2000-DSPE: TopFluor-Chol at 96.5:3:0.5 mol % dissolved in chloroform at a concentration of 3.5 mg/mL. Working in a chemical safety hood, we deposit 2 mL of the lipid solution using the L-shaped glass delivery tube similar to how the dextran solution was coated onto the paper. The lipid-coated PCP was placed into a standard laboratory vacuum chamber for 1 hour to remove traces of organic solvent. We then place the lipid-coated PCP in a 330 mm × 279 mm (13-in. × 11-in.) baking tray (inner dimension, 290 mm × 188 mm (11.4-in × 7.4-in)) with PTFE magnetic stir bars placed on the corners of the paper to prevent curling of the paper upon hydration. The paper was hydrated with 200 mL of 1× PBS + 100 mM sucrose. We incubate the sample for 1 hour before removing the stir bars. We harvest the GUVs using a flexible shrink film (Grafix art) cut to a dimension of 188 mm × 127 mm (7.4-in. × 5-in). We move the film across the 279 mm (11-in.) edge of the PCP while applying gentle pressure to detach the GUV buds from the surface. We repeat the process a second time. We transfer the solution containing the GUVs into four 50 mL Falcon tubes for storage. Quantification of GUV yield was performed similarly to samples prepared in 48-well plates.

### Preparation of a tissue-like close-packed layer of GUVs

To obtain the image shown in Figure 5b, we mix 7.5 mL of the sample with 7.5 mL of 1×PBS+100 mM glucose in a 15 mL Falcon tube and centrifuge at 2080× g for 30 minutes to pellet the GUVs.

We then remove 14 mL of the supernatant. Next, using a micropipette, we take a 10 µL aliquot from the bottom of the remaining 1 mL and mix the aliquot with 50 µL of 1× PBS+100 mM glucose and place the suspension into an imaging chamber. We allowed the GUVs to sediment for 3 hours before imaging. We acquired confocal Z-stacks starting from the bottom of the chamber and extending to a height of 35.8 µm (47 slices, 425.1 µm × 425.1 µm, 3568 pixels × 3568 pixels, with a slice thickness of 1.52 µm and slice interval 0.762 µm) using a 20× 1.0 NA Plan-Apochromat objective with a pinhole of 1.02 A.U.

### Characterizing the long-term functionality of PCP

Two separate pieces of ULGT agarose PCP were stored at 4 °C and 22 °C in 150 mm polystyrene Petri dishes. We assemble GUVs from both stored pieces of ULGT agarose PCP and measure the molar yields at 1, 4, 7, 14, 21 and 28 days. We additionally assemble GUVs from a piece of ULGT agarose PCP stored at 4 °C for 28 days without removing it from the low temperature incubator.

## Author contributions

A.B.S. conceived and directed the study. A.C. performed experiments and analyzed the data. V.V. performed the scale up experiments. J.P. performed preliminary experiments. A.B.S and A.C. wrote the manuscript. All authors have given approval to the final version of the manuscript.

## Data availability

The data supporting this article have been included in the manuscript.

## Acknowledgements

This work was funded by the National Science Foundation through NSF CAREER DMR-1848573 and NSF-CREST: Center for Cellular and Biomolecular Machines at the University of California, Merced, NSF-HRD-1547848 and NSF-EES-2112675. The data in this work was collected, in part, with a confocal microscope acquired through the National Science Foundation MRI Award Number DMR-1625733, and a scanning electron microscope acquired through NASA Grant NNX15AQ01A that are housed and managed by the Imaging and Microscopy Core at UC Merced. We would like to thank Brad Bartholomai Ph.D. and our local team from Nikon Instruments for their assistance in obtaining the superresolution images of actin in GUVs. We would like to thank Kennedy Nguyen for help with SEM imaging. The schematic in Figure 6 was created in BioRender. Cooper, A. (2025) https://BioRender.com/g69f517.

