## Supporting Information for "Combining Nanoscale Curvature and Polymer Osmotic Pressure for Efficient Giant Vesicle Assembly under Physiological Conditions"

### Supporting Text

#### Estimating the range of surface concentrations of polymers used in the literature

There are several methods in the literature to prepare films of agarose and polyvinyl alcohol (PVA) on glass surfaces. We calculate the nominal surface concentration of the polymer, polymer-NSC, in the literature using Equation S1.

$$\text{Polymer} - \text{NSC} = \frac{w_{ps}V_{ps}}{M_p A_{substrate}} \qquad \text{Equation S1}$$

In this equation, $w_{ps}$ is the concentration of the polymer solution (g/mL), $V_{ps}$ is the volume of the polymer solution deposited onto the substrate (mL), $M_p$ is the molar mass of the polymer (g/mol), and $A_{substrate}$ is the area of the substrate ($cm^2$). We assume a density, ρ = 1 g/mL for unit conversion from % w/w to g/mL. We restrict our analysis to assembly of GUVs using the gel-assisted method in salty solutions and do not consider reports of gel-assisted assembly in low salt solutions.

Reference[1] fabricates films of ultralow gelling temperature (ULGT) agarose on glass slides (75 × 50 × 1 mm, Corning Glass Works, Corning, NY) using two different methods, 1) by dip coating the slides in a remelted solution of 1 % w/w agarose in deionized water or, 2) by depositing ~ 300 µL of a remelted solution of 1 % w/w agarose in deionized water on the surface before spreading with the side of a pipette tip. The glass slides with the agarose solution were placed on a hot plate kept at 40 ºC and allowed to dry for 1 – 3 hrs. We use the values reported for method 2 to calculate polymer-NSC. The molar mass of agarose is 120,000 g/mol.[2]

Reference[3] fabricates films of ultralow gelling temperature (ULGT) on glass coverslips (25 mm × 25 mm, VWR® Micro Cover Glasses, Square, No.1) by depositing ~ 300 µL of 1 %

w/v agarose solution at 65 – 75 ºC on the surface and spreading with the side of a 1000 μL pipette tip. The glass coverslips with the agarose solution were then placed on a Parafilm piece with the polymer side facing up and then into a 37 ºC incubator for at least 1 hr to dehydrate the agarose. Reference[4] uses the same deposition procedure but dehydrates the agarose at 40 ºC for > 1 hour.

Reference[5] fabricates films of ULGT agarose on glass coverslips (25 mm × 25 mm, ChemGlass, Vineland, NJ) by applying 20 μL of a 2 % w/v solution of ULGT agarose at 70 ºC to one coverslip, dropping a second coverslip on top of the first coverslip, and sliding the coverslips apart to create a thin film on both coverslips. We assume a volume of 10 μL of polymer solution per coverslip to calculate polymer-NSC. Reference[6] uses the same procedure to deposit the aqueous polymer solutions and uses ULGT agarose concentrations of 0.1, 0.5, 1.0 3.0, and 5.0 % w/v.

Reference[7] uses 300 μL of 1 % w/w agarose at various gelling temperatures and PVA on square glass coverslips (22 mm × 22 mm, Corning). The coated glass coverslips are then placed on a hotplate set to 40 °C on a piece of Parafilm with the polymer side facing up and allowed to dehydrate for a minimum of 2 hours.

Other references use spin coating to spread the agarose solution on the substrate with no information on the thickness of the aqueous polymer film prior to drying thus preventing the calculation of polymer-NSC.[8,9]

Reference[10] uses 100 - 300 μL of 5 % w/w PVA with a molecular weight of 145,000 g/mol on round glass coverslips (30 mm diameter, Menzel-Gläser). Most authors that use the PVA-assisted hydration protocol use a similar molecular weight, concentration, and volume of the PVA however the sizes of substrates are not reported preventing the calculation of the polymer-NSC.[11–15]

**LGT agarose PCP-assisted hydration**

We previously found that high yields of GUVs can be obtained only when partially soluble LGT agarose was used and only when assembly was performed at room temperature (22 ºC) when glass was used as a substrate, i.e. LGT agarose PCG.[7] The main text focused on reporting results with ULGT agarose since it showed wider compatibility with assembly at different temperatures. Here we investigate the behavior of LGT agarose on nanocellulose paper at various temperatures and incubation times to further understand the behavior of LGT agarose PCP for assembling GUVs in salty solutions. We found that the optimal polymer-NSC of LGT agarose was 1.5 nmol/cm$^2$ on nanocellulose paper.

We evaluated the effect of incubating for 2 hours as performed in reference.[7] We perform PCP-assisted hydration by incubating for 1 hour in this work. The yield at 2 hours, 16 ± 1.5 % (Table S2), was not significantly different from the previously reported yield on LGT agarose-coated coverslips which was 17 ± 1 % ($p$ = 0.497). However, the yield at 1 hour was significantly lower at 15 ± 0.5 % ($p$ = 0.0245) (Table S2). We conclude that for the partially soluble LGT agarose, a long incubation time is necessary for maximal yield. In comparison, using ULGT agarose, hyaluronic acid, and dextran PCP resulted in GUV yields of 21 ± 1 %, 15 ± 2 %, and 15 ± 0.3% at 1 hour respectively (Table 1).

Consistent with the observation that LGT agarose is partially soluble, at 4 ºC, using LGT agarose PCP as a substrate resulted in a GUV yield of 9 ± 1 % (Table S2). We increased the incubation time to 24 hours to test if a significantly increased incubation time could result in a high yield of GUVs. We chose an incubation time of 24 hours since we expect that biomolecular degradative processes and the rate of growth of microorganisms to be lower at 4 ºC compared to 22 ºC. We found that the yield increased to 13 ± 0.5 % ($p = 0.004$). Despite the long incubation time, similar to our results at 22 ºC, the yield of GUVs that we obtained using LGT agarose PCP was lower than the yield of GUVs that we obtained using ULGT agarose, hyaluronic acid, and dextran PCP at 1 hour, which were 22 ± 2 %, 20 ± 1 %, and 18 ± 1% respectively.

We also evaluated the yield of GUVs obtained using LGT agarose PCP at 37 ºC. This temperature is above the gelling temperature of LGT agarose. The increased solubility of LGT agarose at this temperature resulted in dewetting of the agarose and significantly lowered yields of GUVs.[7] The yield of GUVs obtained from LGT agarose PCP substrate was 18 ± 3 % after 1 hour of incubation (Table S2). This yield was significantly higher than LGT agarose-coated coverslips at a polymer-NSC of 5.2 nmol/cm$^2$ and PCG at a polymer-NSC of 1.5 nmol/cm$^2$ which had yields of 2.2 ± 0.1 % [7] and 5.0 ± 0.7 % respectively (Table S3). Thus, akin to our observations for the other soluble polymers on PCP, nanocellulose paper is effective at limiting the scale of dewetting patterns on LGT agarose-PCP at temperatures where LGT agarose becomes soluble.

We conclude that the use of ULGT agarose, dextran, and hyaluronic acid PCP at their respective optimal polymer-NSCs allows the widest range of temperature for high yields at a short incubation time of 1 hour. However, these soluble polymers have low yields when glass is used as a substrate due to dewetting (Figure S8). If use of glass is essential, then only LGT agarose PCG and only at temperatures where LGT agarose is partially soluble (~ 22 ºC) and an incubation time of 2 hours results in comparable yields to PCP.

**Supporting Figures**

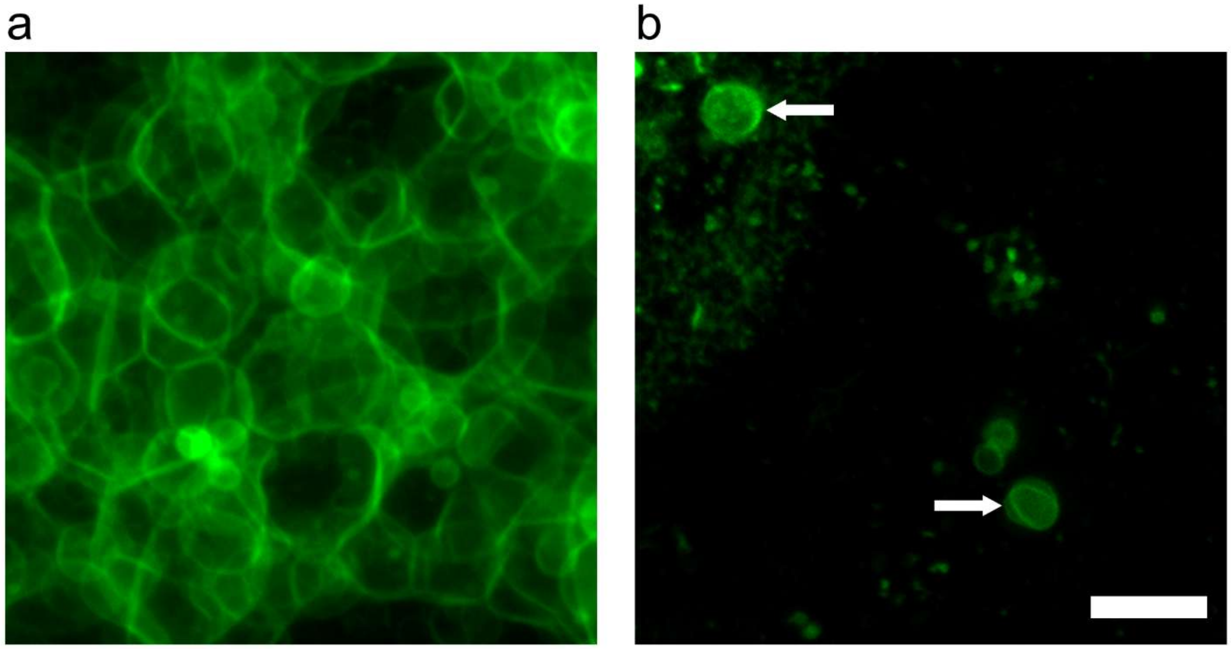


**Figure S1.** ULGT agarose-coated PCP before and after harvesting. a) Summed z-stack of 100 µm from the surface of the substrate before harvesting. b) summed z-stack of 100 µm from the surface of the substrate after harvesting. White arrows point to free-floating, detached GUVs.

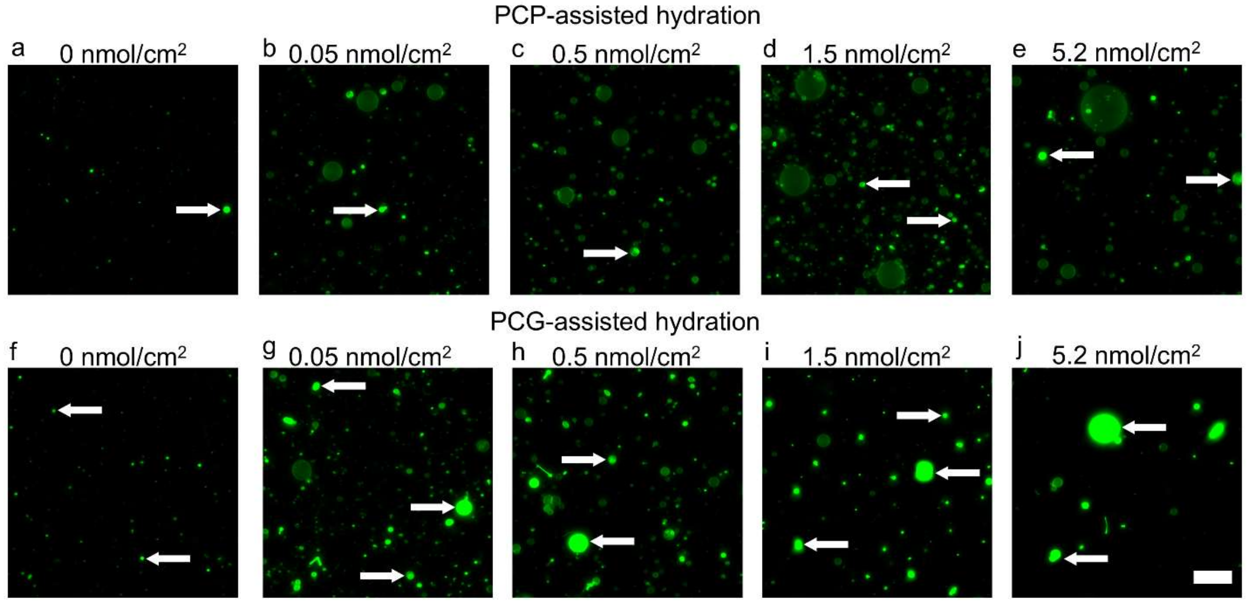


**Figure S2.** Representative confocal fluorescence microscope images of the harvested objects from the samples shown in Figure 1. The white arrows highlight non-GUV structures. Scale bar is 50 μm.

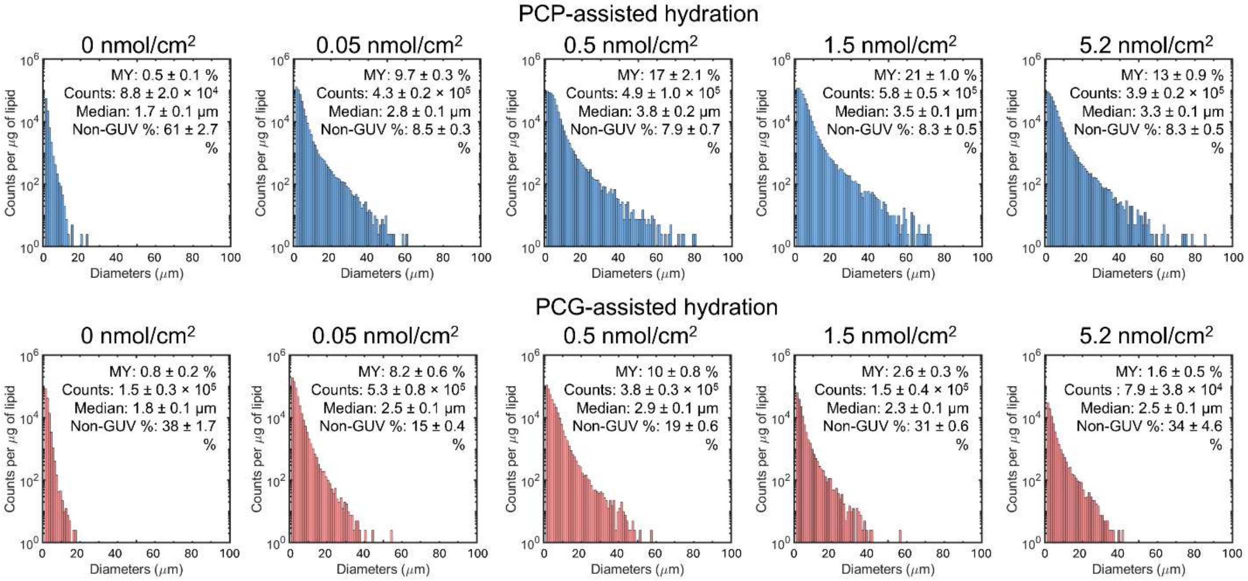


**Figure S3.** Histogram of size distribution of GUVs from Figure 1 in the main text. Each histogram is the average of N=3 independent repeats. Note the logarithmic scale on the *y*-axis. Bin widths are 1 μm. The molar yield (MY), total counts per μg of lipid (counts/μg), median diameter and non-GUV % are reported in the text inserts. The values are reported as the mean ± 1 standard deviation.

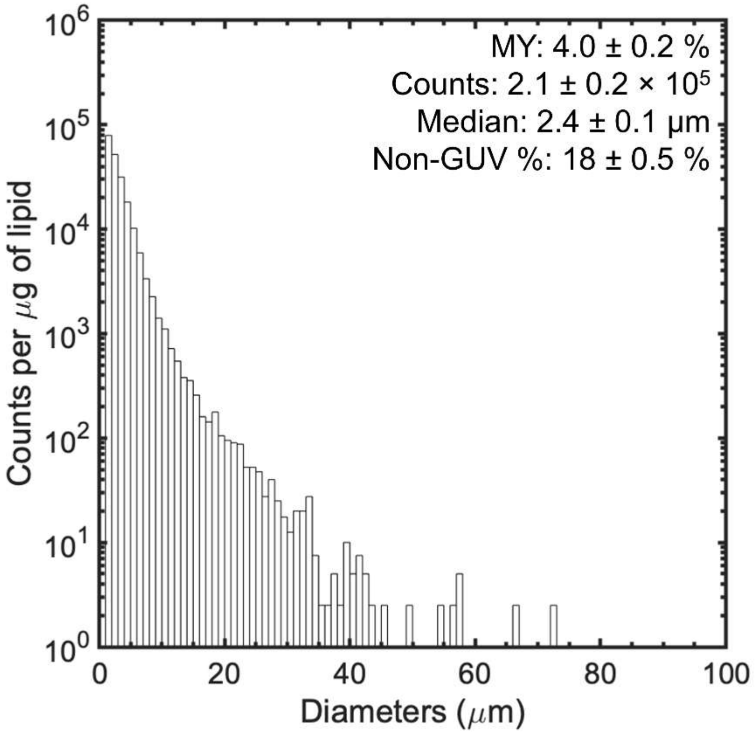


**Figure S4.** Histogram of size distribution of GUVs from assembly on frosted glass slides as shown in Figure 3 in the main text. Note the logarithmic scale on the y-axis. Bin widths are 1 μm. The molar yield (MY), total counts per μg of lipid (counts/μg), median diameter, and non-GUV % are reported in the text inserts.

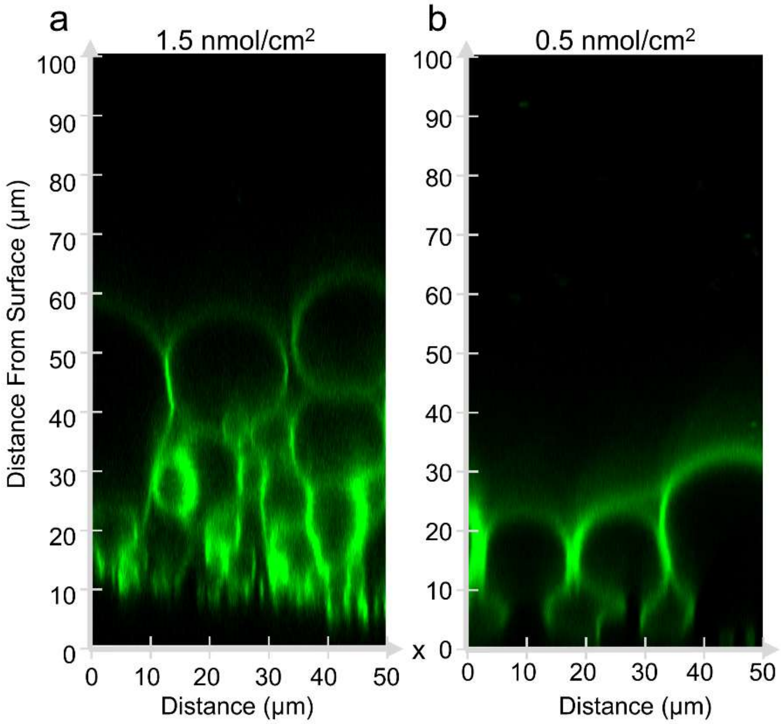


**Figure S5.** Orthogonal x-z images without histogram equalization shown in Figure 4 in the main text.

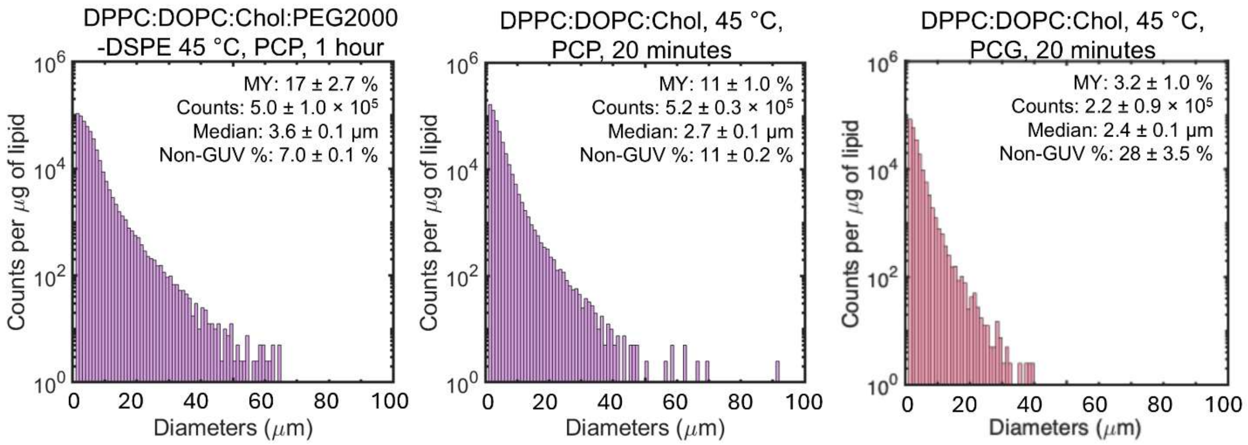


**Figure S6.** Histogram of size distribution of GUVs from Figure 5a,d in the main text. Each histogram is the average of N=3 independent repeats. Note the logarithmic scale on the *y*-axis. Bin widths are 1 µm. The molar yield (MY), total counts per µg of lipid (counts/µg), median diameter, and non-GUV % are reported in the text inserts. The values are reported as the mean ± 1 standard deviation.

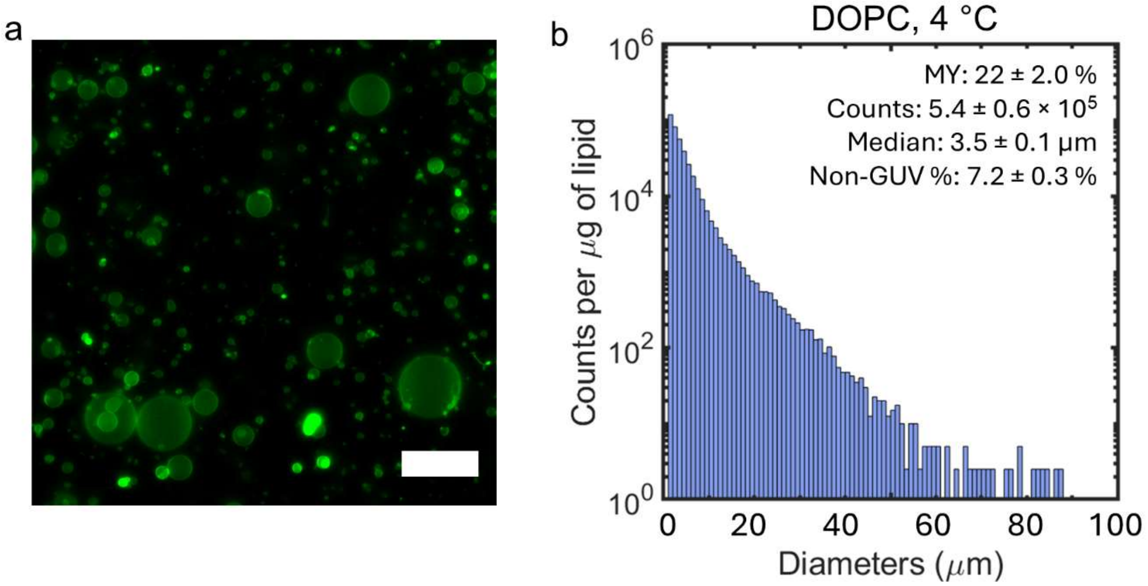


**Figure S7.** a) Representative harvested image of DOPC + 3 mol % PEG2000-DSPE assembled at 4 ºC from Figure 5e. Scale bar is 50 µm. b) The histogram is the average of N=3 independent repeats. Note the logarithmic scale on the *y*-axis. Bin widths are 1 µm. The molar yield (MY), total counts per µg of lipid (counts/µg), median diameter, and non-GUV % are reported in the text inserts. The values are reported as the mean ± 1 standard deviation.

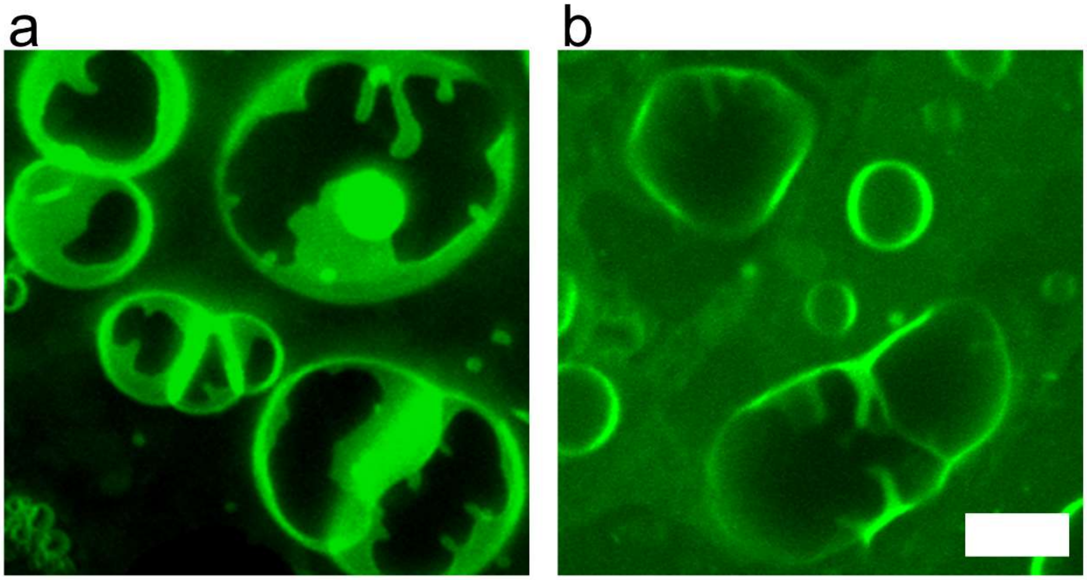

**Figure S8.** Hyaluronic acid and dextran show significant dewetting when glass slides are used as a supporting substrate. a) 0.05 nmol/cm$^2$ % hyaluronic acid 1,250-1,500 kDa. b) 0.31 nmol/cm$^2$ dextran 100 kDa. Scale bar is 15 μm**.**

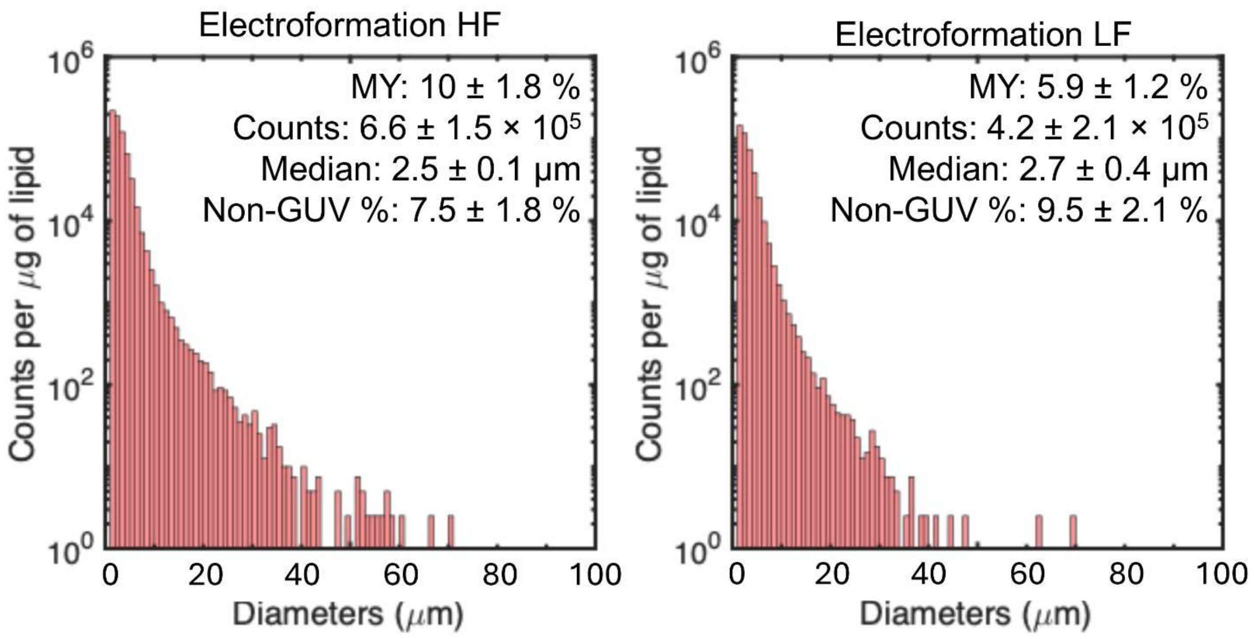


**Figure S9.** Histogram of size distribution of GUVs from Figure 6 in the main text. Each histogram is the average of N=3 independent repeats. Note the logarithmic scale on the *y*-axis. Bin widths are 1 µm. The molar yield (MY), total counts per µg of lipid (counts/µg), median diameter, and non-GUV % are reported in the text inserts. The values are reported as the mean ± 1 standard deviation.

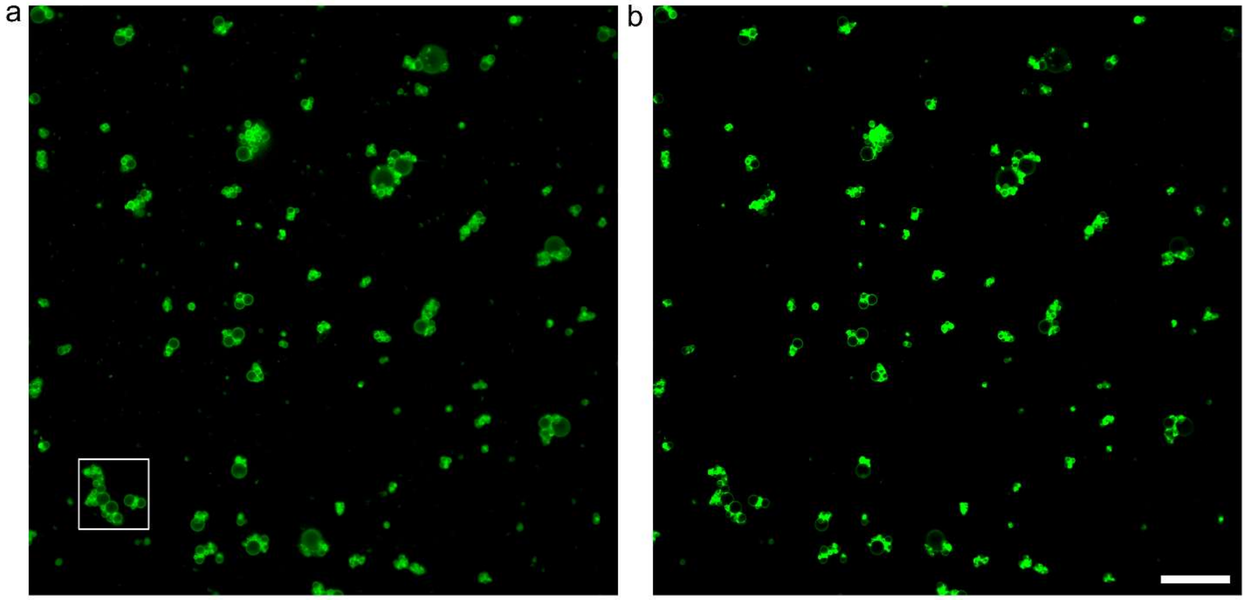


**Figure S10.** a) Full frame image (425.1 μm × 425.1 μm) illustrating the clusters of GUVs that are obtained when the membrane contained only DOPC. The histogram of the image has been equalized to show both bright and dark objects within an 8-bit dynamic range. The white box is the cropped cluster show in Fig. 7b in the main text. b) The image without histogram equalization. The scale bar is 50 μm.

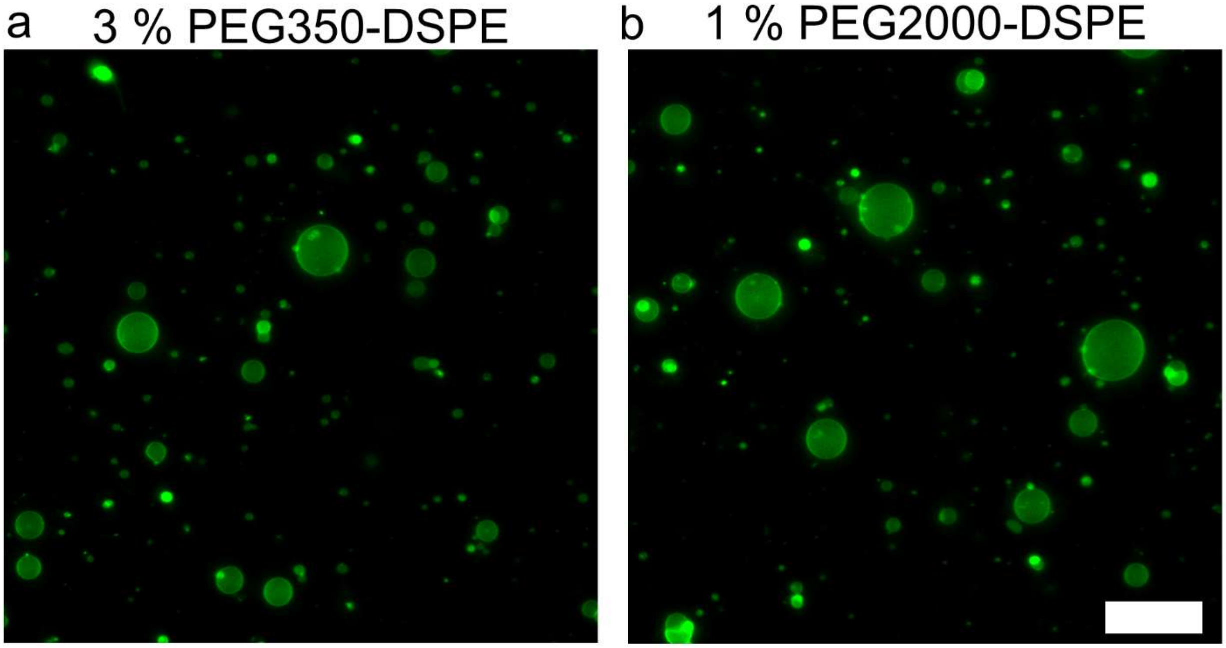


**Figure S11.** Representative harvested image of a) DOPC + 3 mol % PEG350-DSPE and b) 1 mol % PEG2000-DSPE assembled on 1.5 nmol/cm$^2$ ULGT agarose PCP from Figure 7d,e. Scale bar is 50 μm.

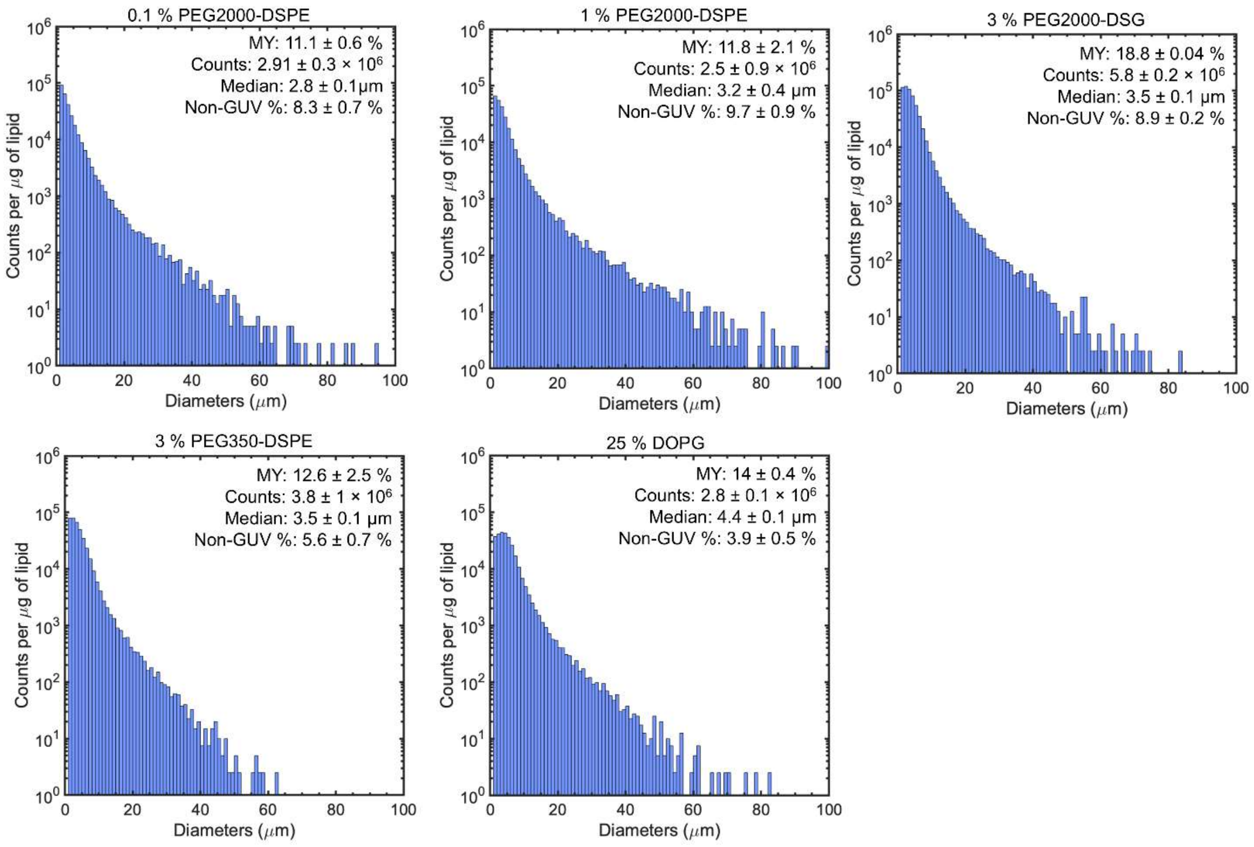


**Figure S12.** Histogram of size distribution of GUVs from Figure 7d,e in the main text. Each histogram is the average of N=3 independent repeats. Note the logarithmic scale on the *y*-axis. Bin widths are 1 μm. The molar yield (MY), total counts per μg of lipid (counts/μg), median diameter, and non-GUV % are reported in the text inserts. The values are reported as the mean ± 1 standard deviation.

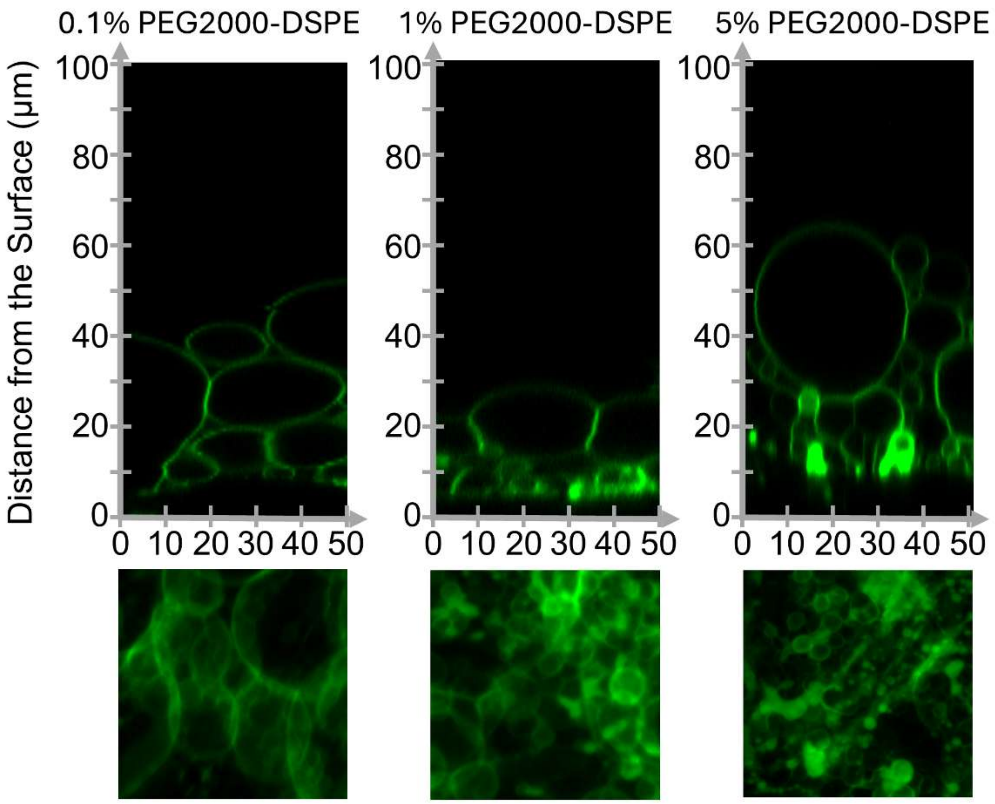


**Figure S13.** Configuration of hydrated DOPC films with different mol % of PEG2000-DSPE. The upper panels are orthogonal reconstructions. The lower panels are summed z-projections of the first 19 µm from the surface.

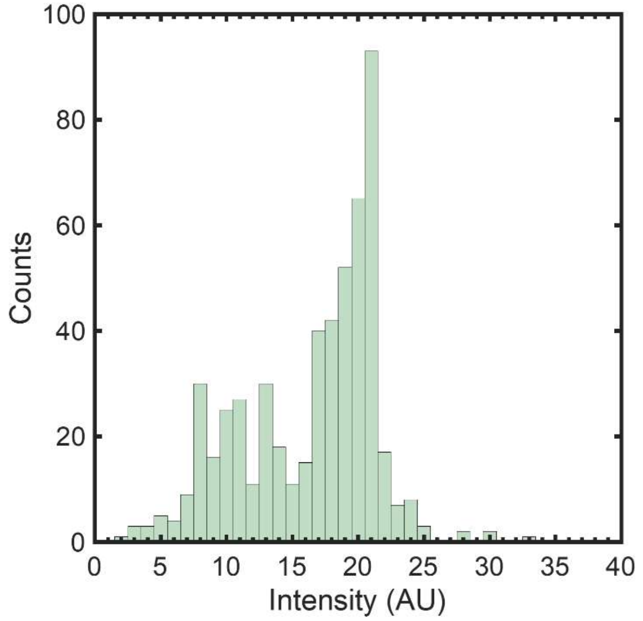


**Figure S14.** Histograms of luminal intensity of GFP from the second independent sample of the myTXTL reaction in GUVs. N= 540 GUVs. Mean intensity 16 ± 5, lowest intensity 2.5, highest intensity 33, CV =0.31. We estimate there were 1.15 × $10^5$ GUVs that expressed deGFP in the chamber.

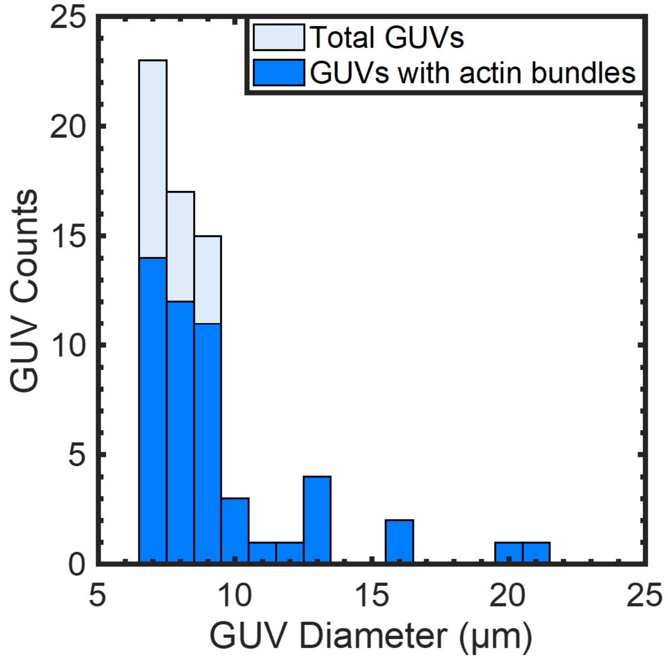


**Figure S15.** Fraction of actin bundles seen in GUVs vs the diameter of GUVs. n=68 GUVs. The estimated number of GUVs in the chamber with actin bundles with diameters > 6.5 µm is $1.1 \times 10^5$.

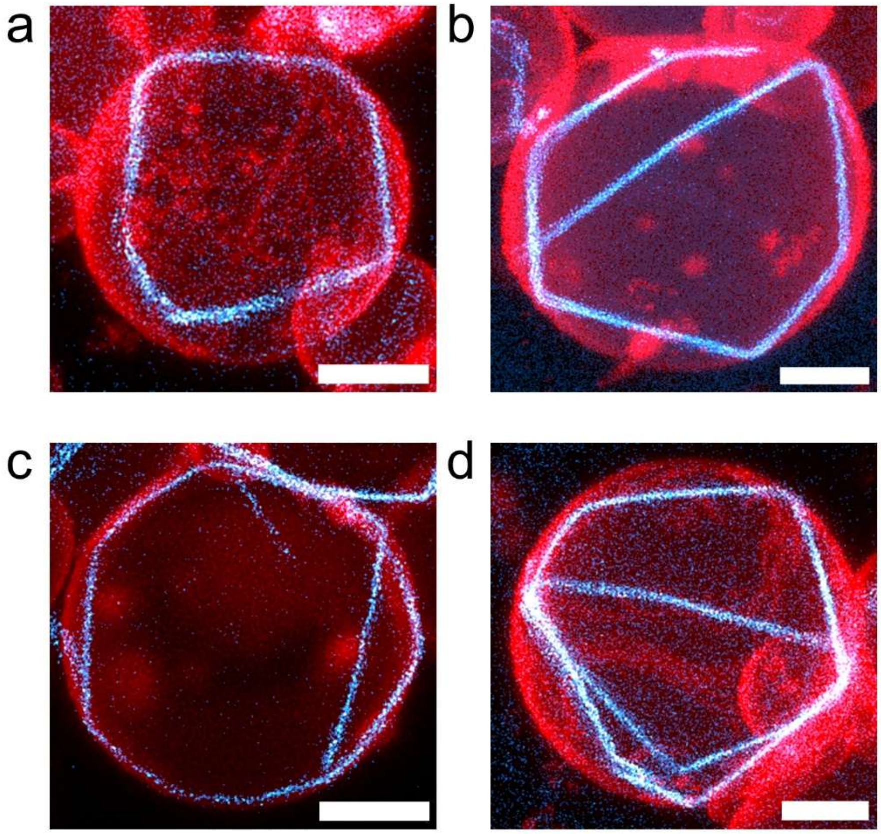


**Figure S16.** Two-channel superresolution images of actin ring conformations within GUVs prepared through PCP-assisted hydration. The images are an overlay of the membrane channel, false colored red, and the actin channel, false colored cyan. The concentration of actin and fascin used is 12 μM and 1.8 μM fascin respectively. a) A GUV with a single actin ring. b) A GUV with a single actin ring that did not connect. c) A GUV with a single actin ring with one branch. d) A GUV with multiple actin rings. Scale bars are 5 μm.

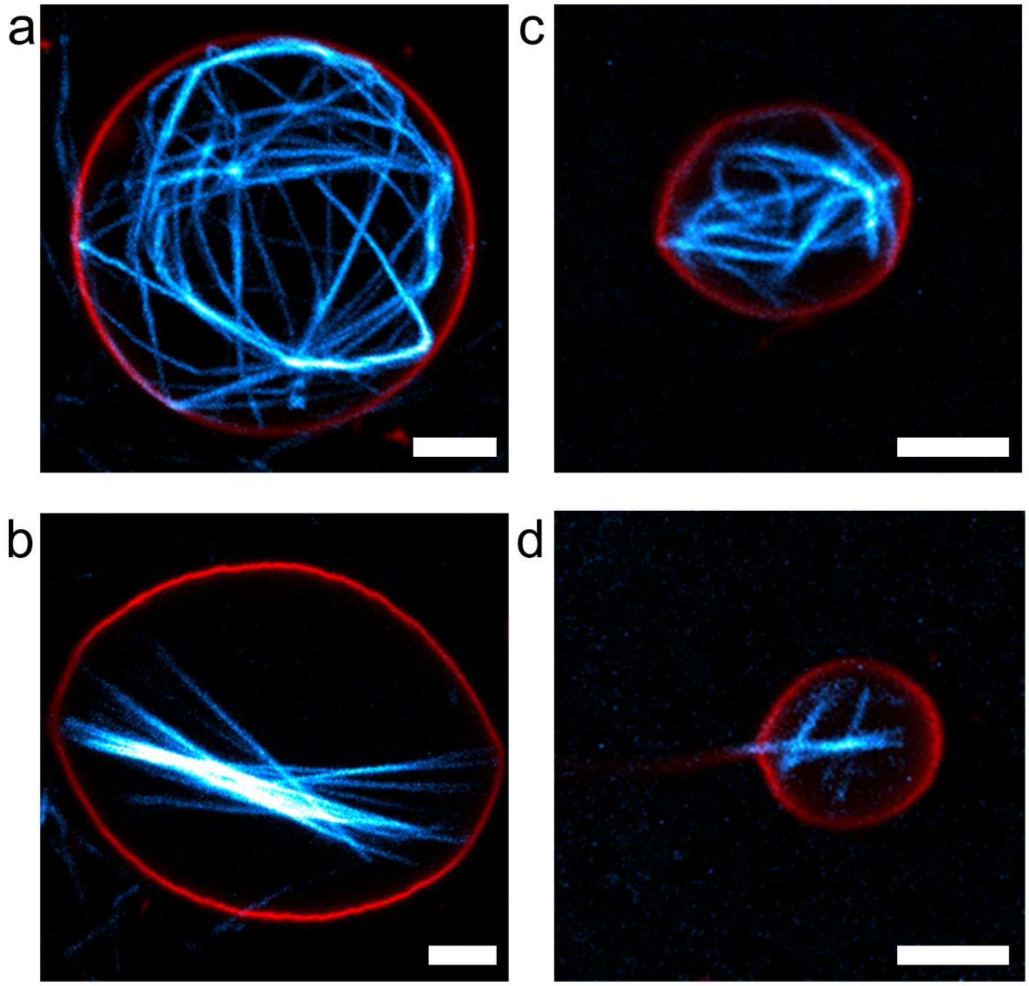


**Figure S17.** Images of GUVs with multiple actin bundles and actin protrusions assembled with high concentrations of actin. The images are an overlay of the membrane channel, false colored red, and the actin channel, false colored cyan a-b) 24 µM actin and 1.5 µM fascin and c-d) 50 µM actin and 3 µM fascin. Scale bars are 5 µm.

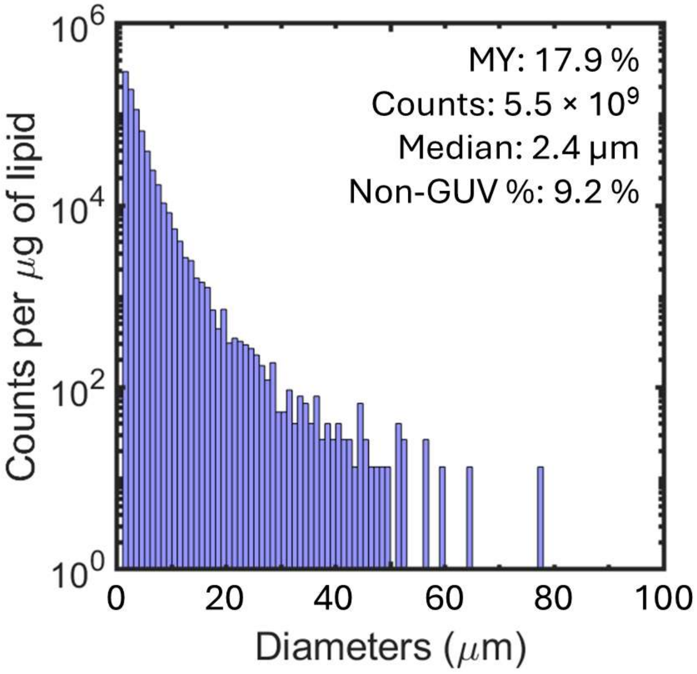


**Figure S18.** Histogram of size distribution of GUVs from large scale assembly of GUVs shown in Figure 9 in the main text. Note the logarithmic scale on the y-axis. Bin widths are 1 μm. The molar yield (MY), total counts per μg of lipid (counts/μg), median diameter, and non-GUV % are reported in the text inserts.

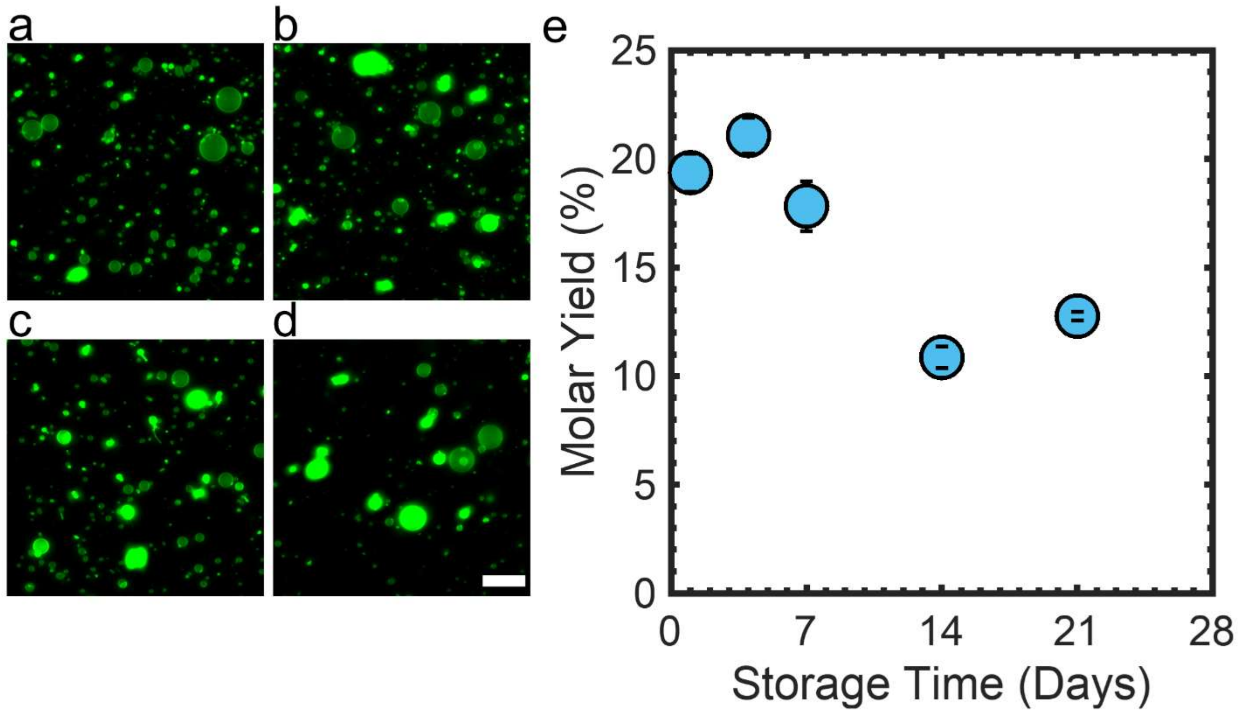


**Figure S19.** Repeated removal of ULGT agarose PCP from the fridge results in rapidly decreasing yield. Representative images of objects harvested from ULGT agarose PCP that was stored at 4 °C and repeatedly removed from low temperature storage to perform PCP-assisted hydration. The PCP was removed once at each timepoint, three 9.5 mm diameter disks punched out and then placed back into the fridge. a) 4 days, b) 7 days, c) 14 days and d) 21 days. e) Plot of the evolution of molar yield with storage time. The points are average of 3 independent repeats and the error bars are one standard deviation from the mean. Scale bar is 50 μm.

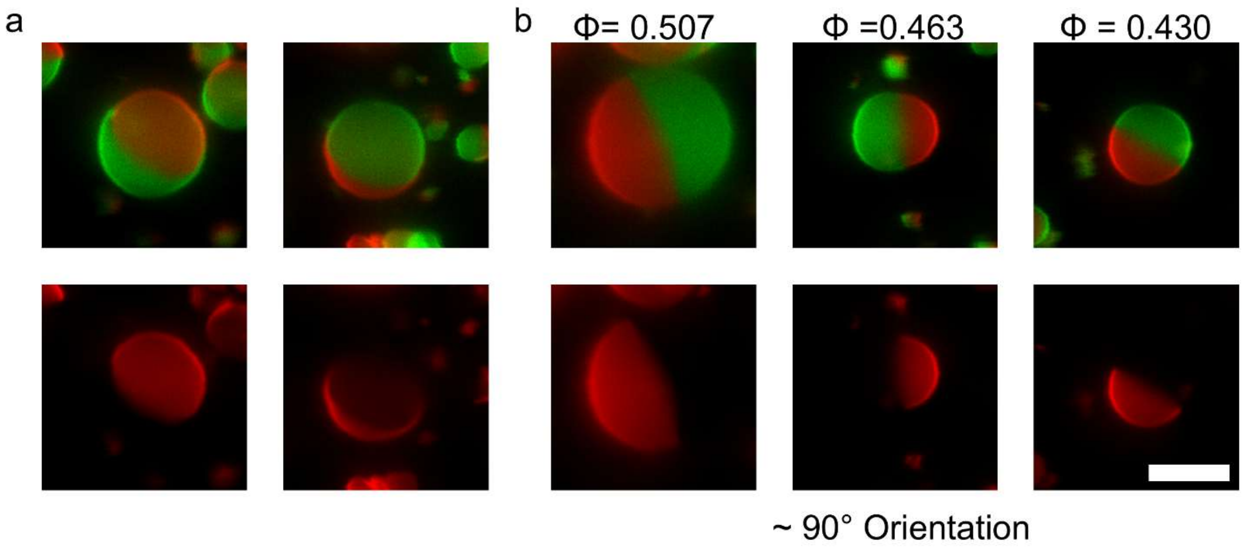


**Figure S20.** Representative images of phase-separated GUVs. The DOPC rich $L_d$ phase is false colored red. The DPPC and Cholesterol rich $L_o$ phase is false colored green. GUVs sediment in random orientations in the chamber. a) Examples of GUVs that do not appear oriented at a 90° angle to the imaging plane. b) Examples of GUVs that appear oriented at a 90° angle to the imaging plane. $\phi$ is the fraction of the $L_d$ phase. Scale bar is 10 μm.

### Supporting Tables

**Table S1.** Polymer type, molecular weight, substrate, and polymer-NSC assembly used in literature.

| Polymer type | Molecular Weight (g/mol) | Substrate | Polymer-NSC (nmol/cm$^2$) | Reference |
|---|---|---|---|---|
| Agarose, ULGT | 120,000 | Glass slide | 0.67 | [1] |
| Agarose, ULGT | 120,000 | Glass coverslip | 0.27 | [5] |
| Agarose, ULGT | 120,000 | Glass coverslip | 4.0 | [3,4] |
| PVA | 145,000 | Glass coverslip | 4.89 – 14.64 | [10] |
| Agarose, LGT | 120,000 | Glass coverslip | 0.01 – 0.67 | [6] |
| Agarose, LGT* | 120,000 | Glass coverslip | 5.2 | [7] |
| PVA** | 146,000-186,000 | Glass coverslip | 4.25 | [7] |

* Authors also used ULGT agarose, Medium Gelling Temperature (MGT) agarose, and High Gelling Temperature (HGT) agarose

** We use the low end of molecular weight range to calculate polymer-NSC.

**Table S2:** Molar yields, counts per µg, and percent non-GUV objects from LGT agarose PCP.

Each value is provided as mean ± standard deviation of N=3 independent repeats.

| Polymer | Polymer-NSC (nmol/cm$^2$) | Lipid Composition | Molar Yield (%) | GUV Counts ($\times 10^5$ GUVs/µg) | Percent non-GUVs (%) |
|---|---|---|---|---|---|
| **4°C** | | | | | |
| Agarose, LGT 120 kDa | 1.5 | DOPC:PEG 2000-DSPE | 8.6 ± 1.0 | 5.4 ± 1.2 | 8.4 ± 0.4 |
| Agarose, LGT, 120 kDa, 24-hour hydration | 1.5 | DOPC:PEG 2000-DSPE | 13.3 ± 0.5 | 6.5 ± 0.7 | 7.0 ± 0.2 |
| **22 °C** | | | | | |
| Agarose, LGT 120 kDa | 1.5 | DOPC:PEG 2000-DSPE | 14.8 ± 0.5 | 10.5 ± 0.3 | 4.7 ± 0.1 |
| Agarose, LGT, 120 kDa, 2-hour hydration | 1.5 | DOPC:PEG 2000-DSPE | 16.2 ± 1.6 | 12.7 ± 1.2 | 6.7 ± 0.5 |
| **37°C** | | | | | |
| Agarose, LGT 120 kDa | 1.5 | DOPC:PEG 2000-DSPE | 18.3 ± 2.6 | 5.4 ± 1.0 | 7.5 ± 0.3 |
| **45°C** | | | | | |
| Agarose, LGT 120 kDa | 1.5 | DOPC:DPPC :Chol:PEG 2000-DSPE | 15.1 ± 1.8 | 4.8 ± 0.6 | 8.0 ± 1.2 |

**Table S3:** Miscellaneous molar yields, counts per μg, and percent non-GUV objects from various PCPs. The NSC-polymer has not been optimized in this table. Each value is provided as mean ± standard deviation of N=3 independent repeats. When less than three repeats were performed, the raw numbers are provided.

| Polymer | Polymer-NSC (nmol/$cm^2$) | Lipid Composition | Molar Yield (%) | GUV Counts ($\times 10^5$ GUVs/μg) | Percent non-GUVs (%) |
|---|---|---|---|---|---|
| **4°C** | | | | | |
| Hyaluronic Acid, 8-15 kDa | 2.9 | DOPC:PEG 2000-DSPE | 7.6 ± 1.9 | 4.1 ± 1.5 | 8.9 ± 1.1 |
| **22 °C** | | | | | |
| Dextran, 100 kDa | 1.9 | DOPC:PEG 2000-DSPE | 11.6 ± 0.6 | 4.0 ± 0.1 | 11.7 ± 2.2 |
| Dextran, 6 kDa | 103 | DOPC:PEG 2000-DSPE | 4.7 | 2.0 | 2.7 |
| Dextran, 6 kDa | 52 | DOPC:PEG 2000-DSPE | 7.3, 6.2 | 2.2, 1.9 | 5.5, 5.0 |
| DNA, Single Stranded from Salmon Testes | 4.6 | DOPC:PEG 2000-DSPE | 1.4, 1.5 | 1.0, 1.3 | 11, 11 |
| Hyaluronic Acid, 8-15 kDa | 2.9 | DOPC:PEG 2000-DSPE | 8.5 ± 1.7 | 4.5 ± 1.0 | 9.0 ± 0.5 |
| PVA, 31-50 kDa | 6.0 | DOPC:PEG 2000-DSPE | 14.0 ± 1.0 | 6.5 ± 0.5 | 11.2 ± 0.1 |
| **45°C** | | | | | |
| Agarose, ULGT 120 kDa | 0.5 | DOPC:DPPC :Chol:PEG 2000-DSPE | 11.9 ± 1.5 | 3.5 ± 0.9 | 16.2 ± 2.2 |
| PVA, 146-186 kDa | 5.3 | DOPC:DPPC: Chol:PEG 2000-DSPE | 8.4 ± 1.2 | 2.8 ± 0.7 | 18.0 ± 1.9 |

**Table S4.** Molar yield, counts per μg, and percent non-GUV objects from the PAPYRUS method on bare nanocellulose paper in 1×PBS + 100 mM sucrose. Each value is a mean ± standard deviation of N=3 independent repeats.

| Substrate | Polymer-NSC (nmol/cm$^2$) | Lipid Composition | Molar Yield (%) | GUV Counts (×10$^5$ GUVs/μg) | Percent non-GUVs (%) |
|---|---|---|---|---|---|
| **4°C** Bare Nanocellulose Paper | - | DOPC:PEG 2000-DSPE | 0.5 ± 0.1 | 1.9 ± 0.2 | 50.3 ± 0.7 |
| **22°C** Bare Nanocellulose Paper* | - | DOPC:PEG 2000-DSPE | 0.5 ± 0.1 | 3.3 ± 0.8 | 61.4 ± 2.6 |
| **37°C** Bare Nanocellulose Paper | - | DOPC:PEG 2000-DSPE | 1.0 ± 0.2 | 1.8 ± 0.3 | 63.7 ± 1.5 |
| **45°C** Bare Nanocellulose Paper | - | DOPC:DPPC :Chol:PEG 2000-DSPE | 2.4 ± 0.3 | 2.3 ± 0.8 | 27.1 ± 0.2 |

* Data is from the sample plotted in Figure 1a and reproduced here for ease of comparison in tabular format.

**Table S5.** Miscellaneous molar yields, counts per µg, and percent non-GUV objects from various PCGs. The NSC-polymer has not been optimized in this table. Each value is the mean ± standard deviation of N=3 independent repeats.

| Polymer | Polymer-NSC (nmol/cm$^2$) | Lipid Composition | Molar Yield (%) | GUV Counts (×10$^5$ GUVs/µg) | Percent non-GUVs (%) |
|---|---|---|---|---|---|
| **37°C** | | | | | |
| Agarose, LGT, 120 kDa | 1.5 | DOPC:PEG 2000-DSPE | 5.0 ± 0.7 | 2.2 ± 0.5 | 23.8 ± 1.0 |
| **45°C** | | | | | |
| Agarose, ULGT, 120 kDa, 20 minute* hydration | 0.27 | DOPC:DPPC :Chol:PEG 2000-DSPE | 3.2 ± 1.0 | 2.2 ± 0.9 | 28.2 ± 3.5 |
| PVA, 146-186 kDa | 4.3 | DOPC:DPPC: Chol:PEG 2000-DSPE | 13.0 ± 1.7 | 4.2 ± 0.7 | 18.6 ± 1.7 |

* Polymer-NSC chosen to match reference [5]

**Table S6:** ANOVA table of the molar yields of GUVs assembled using different repulsive lipids.

| 'Source' | 'SS' | 'df' | 'MS' | 'F' | 'Prob>F' |
|---|---|---|---|---|---|
| **'Columns'** | 250.618 | 5 | 50.124 | 16.748 | 4.79E-05 |
| **'Error'** | 35.913 | 12 | 2.993 | | |
| **'Total'** | 286.531 | 17 | | | |

**Table S7:** Table of p-values from post hoc Tukey's HSD tests of the molar yields of GUVs assembled using different repulsive lipids. * = $p < 0.05$, ** = $p < 0.01$, *** = $p < 0.001$, NS = not significant.

| Group 1 | Group 2 | *p*-value | Significance | Comments |
|---|---|---|---|---|
| 0.1% PEG2000-DSPE | 1% PEG2000-DSPE | 0.993 | NS | The molar yields between group 1 and group 2 were not significantly different. |
| 0.1% PEG2000-DSPE | 25% DOPG | 0.380 | NS | |
| 0.1% PEG2000-DSPE | 3% PEG350-DSPE | 0.884 | NS | |
| 1% PEG2000-DSPE | 25% DOPG | 0.676 | NS | |
| 1% PEG2000-DSPE | 3% PEG350-DSPE | 0.994 | NS | |
| 3% PEG2000-DSPE | 3% PEG2000-DSG | 0.597 | NS | |
| 25% DOPG | 3% PEG350-DSPE | 0.922 | NS | |
| 25% DOPG | 3% PEG2000-DSG | 0.045 | * | The molar yield from Group 1 was significantly lower than the molar yield from 3% PEG2000-DSG. |
| 0.1% PEG2000-DSPE | 3% PEG2000-DSG | 0.002 | ** | |
| 1% PEG2000-DSPE | 3% PEG2000-DSG | 0.004 | ** | |
| 3% PEG350-DSPE | 3% PEG2000-DSG | 0.009 | ** | |
| 25% DOPG | 3% PEG2000-DSPE | 0.003 | ** | The molar yield from Group 1 was significantly lower than the molar yield from 3% PEG2000-DSPE. |
| 0.1% PEG2000-DSPE | 3% PEG2000-DSPE | 1.44E-04 | *** | |
| 3% PEG350-DSPE | 3% PEG2000-DSPE | 6.68E-04 | *** | |
| 1% PEG2000-DSPE | 3% PEG2000-DSPE | 3.07E-04 | *** | |

**Table S8.** Polymer costs at the ideal surface concentrations.

| Polymer | Optimal polymer-NSC ($nmol/cm^2$) | Cost | Unit of measure | Cost per $cm^2$ at optimal polymer-NSC | Purchase source |
|---|---|---|---|---|---|
| ULGT Agarose | 1.5 | $109.00 | 5 g | $0.0041 | Sigma Aldrich, A2576 |
| Hyaluronic acid 8-15 kDa | 23 | $222.00 | 10 mg | $4.1292 | Sigma Aldrich, 40583 |
| Dextran | 4.3 | $25.90 | 10 g | $0.0011 | Sigma Aldrich, 09184 |
| Carrageenan | 1.2 | $71.10 | 25 g | $0.0018 | Sigma Aldrich, C1013 |
| Poly-D-lysine | 6.2 | $45.32 | 10 mg | $0.9200 | Sigma Aldrich, P0899 |

**Table S9.** Polymer solutions for PCP fabrication.

| Polymer | Concentration (% w/w) | Temperature Solubilized (°C) | Agitation Method | Approximate Time to Dissolution |
|---|---|---|---|---|
| ULGT and LGT agarose | 0.3, 1 | 65 | Vortex every 10 minutes | 30 minutes |
| Bovine serum albumin | 1 | 22 | Pipette half the volume repeatedly until dissolved | 5 minutes |
| Carrageenan | 1 | 65 | Vortex every 10 minutes | 30 minutes |
| Dextran | 1, 4.85 | 22, 65 | Vortex until dissolved for 1 % w/w , vortex every 5 minutes for 4.85 % w/w | 3 minutes for 1 % w/w, ~ 10 minutes for 4.85 % w/w |
| Hyaluronic acid | 1 | 22 | Pipette half the volume repeatedly until dissolved | 5 minutes |
| Poly-D-lysine | 1 | 22 | Pipette half the volume repeatedly until dissolved | 5 minutes |
| Polyvinyl alcohol | 5 | 85 | Vortex every 10 minutes with gradual addition of 1/10$^{th}$ of PVA every 10 minutes until all PVA is added | 3 hours |
| Salmon DNA* | 1 | - | - | - |

*The salmon DNA used was purchased in an aqueous solution at a concentration of 10 mg/mL.
